# SLDP and LIPA mediate lipid droplet-plasma membrane tethering in *Arabidopsis thaliana*

**DOI:** 10.1101/2022.01.13.476213

**Authors:** Hannah Elisa Krawczyk, Siqi Sun, Nathan M. Doner, Qiqi Yan, Magdiel Lim, Patricia Scholz, Philipp Niemeyer, Kerstin Schmitt, Oliver Valerius, Roman Pleskot, Stefan Hillmer, Gerhard H. Braus, Marcel Wiermer, Robert T. Mullen, Till Ischebeck

## Abstract

Membrane contact sites (MCS) are inter-organellar connections that allow for the direct exchange of molecules, such as lipids or Ca^2+^ between organelles, but can also serve to tether organelles at specific locations within cells. Here we identified and characterised three proteins that form a lipid droplet (LD)-plasma membrane (PM) tethering complex in plant cells, namely LD-localised SEED LD PROTEIN (SLDP) 1 and 2 and PM-localised LD-PLASMA MEMBRANE ADAPTOR (LIPA). Using proteomics and different protein-protein interaction assays, we show that both SLDPs associate with LIPA. Disruption of either *SLDP1* and *2* expression, or that of *LIPA*, leads to an aberrant clustering of LDs in Arabidopsis seedlings. Ectopic co-expression of one of the *SLDPs* with *LIPA* on the other hand is sufficient to reconstitute LD-PM tethering in *Nicotiana tabacum* pollen tubes, a cell type characterised by dynamically moving LDs in the cytosolic streaming. Further, confocal laser scanning microscopy revealed both SLDP2.1 and LIPA to be enriched at LD-PM contact sites in seedlings. These and other results suggest that SLDP and LIPA interact to form a tethering complex that anchors a subset of LDs to the PM during post-germinative seedling growth in *Arabidopsis thaliana*.

**One-sentence summary:** SEED LIPID DROPLET PROTEIN1 and 2 and LIPID DROPLET PLASMA MEMBRANE ADAPTOR tether lipid droplets to the plasma membrane in seedlings of *Arabidopsis thaliana*.

## INTRODUCTION

As knowledge on organelle-specific functions and their proteomes has expanded in recent years, there has been ever mounting interest in inter-organelle interactions that are in part facilitated by membrane contact sites (MCS) (Prinz *et al*., 2020). MCS foster physical interactions and the exchange of molecules between organelles without the need of membrane fusion events. The transient connections are established through tethering proteins, connecting the membranes of interacting organelles and allowing for direct exchange of lipids, cellular signals (e.g., Ca^2+^, reactive oxygen species [ROS], etc) and/or other molecules (Baillie *et al*., 2020; Prinz *et al*., 2020; Rossini *et al*., 2020).

It is well recognised that MCS can form between nearly all organelles (Eisenberg-Bord *et al*., 2016; Valm *et al*., 2017; Shai *et al*., 2018; Baillie *et al*., 2020). The endoplasmic reticulum (ER) and peroxisomes, for example, are organelles with well-described interactomes (Shai *et al*., 2016; Zang *et al*., 2020). Also, several multi-organelle MCS have been described, such as the three-way connection between mitochondria, ER, and lipid droplets (LDs) that promotes *de novo* lipogenesis in human adipocytes (Freyre *et al*., 2019).

Although ER-derived LDs are known to engage in various MCS, the LD interactome is less well described than that of other organelles (Bohnert, 2020). LDs consist of a lipophilic core of neutral lipids, such as triacylglycerols (TAGs) and sterol esters, surrounded by a phospholipid monolayer and a variety of surface-associated ‘coat’ proteins. Long believed to be an inert storage organelle, it is now widely accepted that LDs actively participate in a wide range of cellular processes involving lipids and their derivatives (Thiam and Beller, 2017; Welte and Gould, 2017; Ischebeck *et al*., 2020). As such, rather than just housing storage lipids, LDs are now considered to be dynamic hubs for lipid homeostasis and specialised metabolism (Schaffer, 2003). Furthermore, LDs can serve as a sink for reducing cytosolic free fatty acids (Fan *et al*., 2017; Olzmann and Carvalho, 2019; de Vries and Ischebeck, 2020) and ROS (Muliyil *et al*., 2020), and also sequester potentially harmful proteins (Geltinger *et al*., 2020) or histone complexes at the LD surface (Johnson *et al*., 2018).

Given the established role(s) of MCSs in the non-vesicular transport of lipids (Cockcroft and Raghu, 2018), it is not surprising that LDs form contacts with many other organelles (Bohnert, 2020; Valm *et al*., 2017; Gao and Goodman, 2015; Schuldiner and Bohnert, 2017; Rakotonirina-Ricquebourg *et al*., 2021). The majority of these described LD MCS, however, were found mammalian or yeast cells. Knowledge in plants is so far still limited to LD-ER and LD-peroxisome MCS, which are involved in storage lipid accumulation (Cai *et al*., 2015; Greer *et al*., 2020; Pyc *et al*., 2021) and breakdown (Eastmond, 2006; Cui *et al*., 2016), respectively. Likewise, while an LD-plasma membrane (PM) MCS was recently described in fly (*Drosophila melanogaster)* (Ugrankar *et al*., 2019), no such connection has been described for plants.

SEED LD PROTEIN 1 (SLDP1) was lately reported as a plant-specific LD protein (Kretzschmar *et al*., 2020) and, in *Arabidopsis thaliana*, has a close homologue, SLDP2. However, the function(s) of SLDP1 and/or SLDP2 are unknown. We show here that LDs in *sldp1 sldp2* double mutants display an aberrant subcellular positioning (i.e., clustering) during early seedling growth. In proteomic analyses of *sldp* mutants, we identified a novel PM-localised protein that serves as an interaction partner of SLDPs and which we termed LIPA (<u>LI</u>PID DROPLET <u>P</u>LASMA MEMBRANE <u>A</u>DAPTOR). Consistent with this premise, subcellular localisation studies in *Nicotiana tabacum* pollen tubes showed that ectopically-expressed LIPA localises to the PM, but is relocated to apparent PM-LD MCS when co-expressed with SLDP2. Moreover, in both *sldp2* and *sldp1 sldp2* mutants, LIPA is absent from LD-enriched proteome fractions and *lipa* mutants phenocopy *sldp1 sldp2* mutants in terms of the aberrant clustering of LDs in cells during seed germination. Interaction of SLDPs and LIPA was confirmed using fluorescence resonance energy transfer fluorescence lifetime imaging microscopy (FRET-FLIM) and yeast two-hybrid (Y2H) assays. Based on these and other data, we provide a working model, where LIPA anchors SLDP-decorated LDs to the PM in plant cells.

## RESULTS

### SLDP1 and 2 are members of a novel, plant-specific LD protein family and contain a similar LD targeting signal

We recently investigated the proteomes of LD-enriched fractions of Arabidopsis siliques, seeds and seedlings and, in doing so, identified several novel LD protein families (Kretzschmar *et al*., 2020). Notably, some of these proteins were unique to plants, such that they had no obvious homologues in mammals and/or yeast, and they were also annotated to be of unknown function, suggesting that they serve novel roles related to LDs in plants.

One of the families of new Arabidopsis LD proteins we identified with no apparent homology to other proteins and no known function(s) included SLDP1 (AT1G65090) and SLDP2 (AT5G36100). There are three and two splice variants of Arabidopsis SLDP1 and SLDP2, respectively, sharing 35.4% - 46.7% sequence identity (Figure 1A, Supplemental Figure 1). We previously showed that SLDP1.3 targets LDs (Kretzschmar *et al*., 2020) and we confirmed the same LD localisation for SLDP2.1 by transiently expressing the mVenus-tagged protein (i.e., SLDP2.1-mVenus) in *Nicotiana tabacum* pollen tubes (Figure 1B), which pose a well-established model plant cell system for studying LD protein localisation (Müller *et al*., 2017; Müller and Ischebeck, 2018; Kretzschmar *et al*., 2018, 2020).

**Figure 1.**
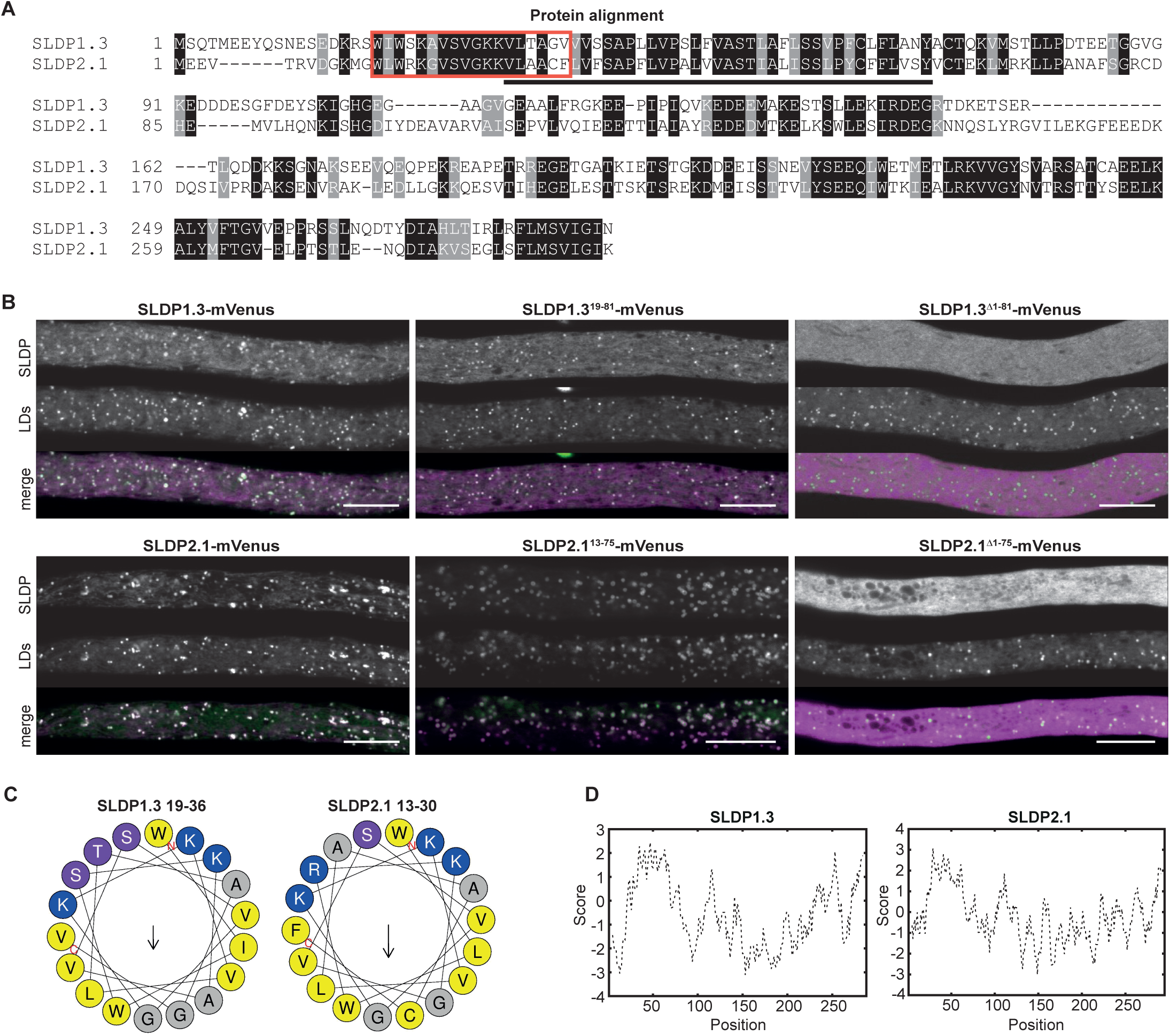
Arabidopsis has two SLDP isoforms that both localise to LDs. **A** Amino acid sequence alignment of SLDP1.3 and SLDP2.1, as generated by T- Coffee. Identical and similar amino acids in SLDP1.3 and SLDP2.1 are shaded black and grey, respectively. Hydrophobic/uncharged regions, as predicted by ExPASy ProtScale, are underlined and potential amphipathic α-helices, as predicted by Heliquest, are boxed. See Supplemental Figure 1 for the protein sequence alignment of five splice variants of Arabidopsis SLDP1 and SLDP2. **B** CLSM images of transiently-expressed full-length and truncated versions of SLDP1.3 and SLDP2.1 appended to mVenus in transiently-transformed *N. tabacum* pollen tubes. Truncated fusion constructs either included sequences predicted to form an amphipathic α-helix and hydrophobic sequences in SLDP1.3 and SLDP2.1 (i.e., SLDP1.3 ^19-81^ and SLDP2.1 ^13-75^) or were devoid of these sequences (i.e., SLDP1.3^Δ1-81^ and SLDP2.1^Δ1-75^); refer also to (A). LDs were stained with Nile red. Images are representative of at least 10 micrographs of transformed pollen tubes per fusion construct. In the merge images, fluorescence attributable to mVenus-tagged SLDP proteins and corresponding Nile red stained LDs are false-colorized magenta and green, respectively; white colour represents co-localisation. Bars, 10 µm. **C** Helical wheel projection of the N-terminal regions of SLDP1.3 (amino acid residues 13-30) and SLDP2.1 (amino acid residues 16-39) predicted by Heliquest to form an amphipathic α-helix. Hydrophobic amino acid residues are coloured yellow, hydrophilic and charged residues are magenta and blue respectively. The direction of the arrow in the helical wheel indicates the position of the hydrophobic face along the axis of the helix. **D** Protein hydropathy profiles of the deduced amino acid sequences of SLDP1.3 and SLDP2.1 based on ProtScale. Note the relatively strong hydrophobic sequence in the N-terminal region of each protein.

As indicated by bioinformatic analyses, both SLDP1.3 and SLDP2.1, as well as their related (spliced) protein variants, share a predicted amphipathic α-helix sequence near their N-termini (amino acids 19-26 and 13-30, respectively) and an adjacent hydrophobic region of ∼40 uncharged amino acids (amino acids residues 31–69 and 25–62, respectively) (Figure 1A, 1C and 1D, Supplemental Figure 2), both of which are known to be key features of the LD targeting signals found in other LD proteins (Kory *et al*., 2016; Olarte *et al*., 2021). To test if these regions are indeed involved in LD targeting of SLDPs, we generated various truncated versions of SLDP1.3 and SLDP 2.1 and expressed them individually in *Nicotiana tabacum* pollen tubes. As shown in Figure 1B, both SLDP1.3^19-81^-mVenus and SLDP2.1^13-75^-mVenus, each including a putative amphipathic α-helix and a hydrophobic sequence, targeted to Nile red-stained LDs, similar to their full-length protein counterparts. By contrast, SLDP1.3^Δ1-81^-mVenus and SLDP2.1^Δ1-75^-mVenus, lacking the corresponding N-terminal portion of each protein, did not target LDs, but instead localised to the cytosol (Figure 1B). Based on these findings, both SLDP1 and 2 proteins appear to share similar LD targeting information.

### Knockout of *SLDP* expression interferes with the subcellular distribution of LDs during post-germinative seedling growth

According to transcriptome data available at Arabidopsis AtGenExpress (Nakabayashi *et al*., 2005; Schmid *et al*., 2005; Winter *et al*., 2007; Waese *et al*., 2017) and the Klepikova eFP browser (Klepikova *et al*., 2016; Waese *et al*., 2017), SLDP1 and SLDP2 are highly expressed in seeds. Consistent with this, we confirmed the expression of all splice variants of SLDP1 and SLDP2 in seeds via two-step reverse transcriptase-quantitative PCR (RT-qPCR) (Supplemental Figure 3). Furthermore, based on proteomic data, both proteins are also found in seedlings (Kretzschmar *et al*., 2020). We therefore reasoned that SLDPs might play a role in LD biology during these stages of development, and we investigated this possibility using a gene loss-of-function approach. Two sets of one Arabidopsis T-DNA and one CRISPR/Cas9 knockout mutant line each for *SLDP1* and *SLDP2* were generated (Figure 2A) and confirmed by genotyping (Supplemental Data S1). In addition, corresponding double knockout mutant lines of *SLDP1* and *SLDP2* were generated. RT-qPCR analyses confirmed that no full-length *SLDP1* or *SDLP2* transcripts were detectable in the *sldp1-1* and *sldp2-1* T-DNA mutant lines, respectively, or in the *sldp1-1 sldp2-1* double mutant (Supplemental Figure 4). *SLDP1* and *SLDP2* transcript levels were not significantly altered in the CRISPR/Cas9 *sldp1-2* and *sldp2- 2* single or double mutant lines (relative to wild-type plants) (Supplemental Figure 4). However, cloning and sequencing of the respective cDNAs from these mutant lines confirmed CRISPR/Cas9-induced mutations (i.e., deletions) that resulted in premature stop codons in the *SLDP1* and *SLDP2* open reading frames (Supplemental Data S1).

**Figure 2.**
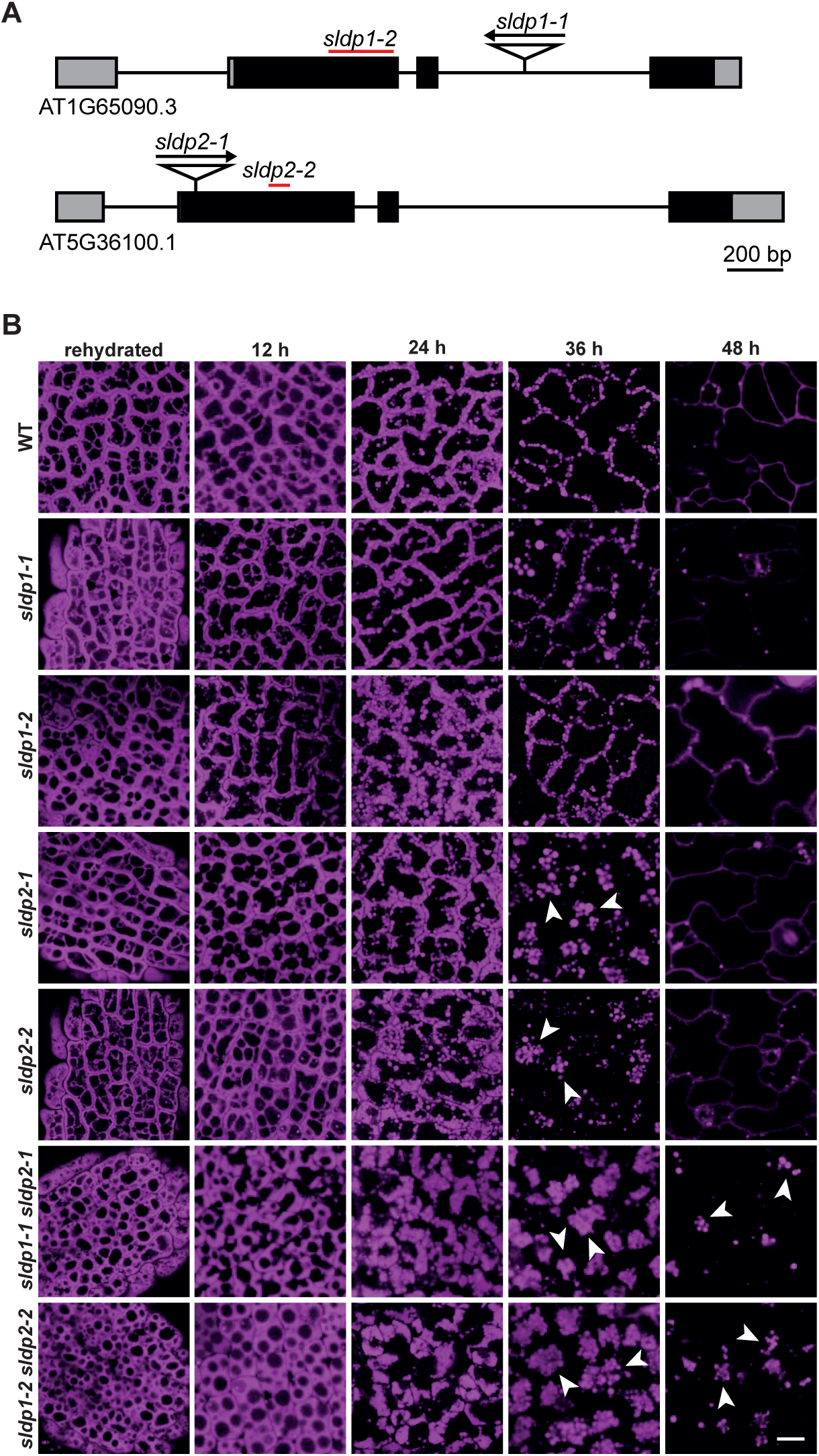
Time-course analysis of LDs in *sldp* mutant Arabidopsis seeds and seedlings. **A** Illustration depicting the Arabidopsis SLDP1.3 and SLDP2.1 genes based on information provided at TAIR. Indicated are the relative positions of the 5’ and 3’ untranslated regions (grey boxes), exons (black boxes), introns (black line), T-DNA insertion sites (triangle, arrow indicating direction of T-DNA), and the regions deleted with CRISPR/Cas9-based genome editing (red lines). **B** CLSM images of rehydrated seeds and seedling cotyledon cells from WT and various *sldp1* and *sldp2* mutant Arabidopsis lines. Seeds were rehydrated for 1 h or stratified for 4 days at 4 °C in the dark. LDs were stained with Nile red after rehydration, or 12, 24, 36 and 48 h (± 2 h) after stratification. Arrowheads indicate obvious examples of LD clusters in *sldp2* single and *sldp1 sldp2* double mutant seedling. Images are single plane images from the middle of the cell (similar planes were chosen for all images). Images are representative of at least five micrographs of seeds and seedlings for each plant line and time point. Bar, 10 µm, applies to all images in the panels.

In comparison to WT plants, neither single nor double T-DNA mutant lines of *SLDP* displayed any obvious growth or developmental phenotypes, including hypocotyl elongation in light- and dark-grown seedlings (Supplemental Figure 5A, B), and were also not strongly affected in the levels of the total fatty acids in seeds (Supplemental Figure 5C). In respect to the 1000 seed weight, the double T-DNA insertion line was also only slightly affected (19.4 ± 0.2 mg in WT and 20.8 ± 0.2 in *sldp1-1* and *sldp2-1*). Furthermore, the degradation of total fatty acids during seed germination and early seeding growth in the *sldp* mutants was not significantly altered compared to WT (Supplemental Figure 5C), including the degradation of eicosenoic acid, which is specifically incorporated into TAGs in Arabidopsis seeds (Rylott *et al*., 2003) (Supplemental Figure 5D). However, an aberrant LD phenotype was readily observed in seedlings from both *sldp2* single and both *sldp1 sldp2* double mutant lines during germination and post-germinative growth. As shown in Figure 2B, and consistent with results presented in other studies of LDs in WT Arabidopsis seeds and seedlings (Cai *et al*., 2015; Gidda *et al*., 2016; Kretzschmar *et al*., 2018), Nile red-stained LDs displayed a ‘typical’ subcellular distribution in rehydrated (mature) seeds and in seedlings from 12 h to 48 h after stratification. That is, in WT seeds and seedlings at 12 h, LDs in cotyledon cells (Figure 2B), as well as those in hypocotyl cells (Supplemental Figure 6), were often closely appressed. At the same stages, no obvious difference in LD morphology or distribution was observed between WT and any of the *sldp* mutant lines (Figure 2B), nor were there any obvious differences in storage vacuole morphology or distribution (Supplemental Figure 7). By 24 h to 48 h, after the completion of germination in Arabidopsis (Bewley, 1997), LDs in cotyledon cells in the WT and both *sldp1* single mutant lines were mostly positioned in close proximity to the PM (Figure 2B). However, at 36 h and 48 h in cotyledon cells of *sldp2* single and particularly *sldp1 sldp2* double mutants, LDs were not evenly distributed along the PM, but instead were noticeably clustered near the centre of the cell. (Figure 2B). A similar clustered LD phenotype was observed in hypocotyl cells in the *sldp1 sldp2* double mutants at 24 h to 48 h, but hypocotyls of the *sldp2* single mutants are less severely affected than its cotyledons (Supplemental Figure 6). Notably, Z-stack images of cotyledon and hypocotyl cells at 36 h, when the LD distribution phenotype was well pronounced (Figure 2B), and subsequent quantification of the LD distribution in these cells, confirmed that a significantly smaller proportion of LDs were located at cell periphery in the *sldp2* single and *sldp1 sldp2* double mutants compared to WT (Supplemental Figure 8). The LD clustering phenotype in cotyledon cells of the *sldp1-1 sldp2-1* double mutant was also observed by electron microscopy, whereas close contacts between LDs and the PM were almost solely observed in WT cotyledon cells (Supplemental Figure 9). Taken together, these data indicate that SLDP1 and SLDP2 are involved in proper subcellular distribution of LDs during post-germinative seedling growth in Arabidopsis.

### Comparative proteomic analyses reveal LIPA as a potential interaction partner of SLDP2

In order to further explore the functions of SLDPs during post-germinative seedling growth in Arabidopsis, we tested if the loss of either SLDP1 or SLDP2, or both proteins, has an influence on the composition of the LD proteome. To this end, proteomic analyses of WT, *sldp1-1*, *sldp2-1* and *sldp1-1 sldp2-1* mutant seedlings (36 h after seed stratification) were performed on LD-enriched fractions, as well as total cellular extracts (abbreviated as TE) (Supplemental Figure 10A-C). Initially, protein levels of SLDP1 and SLDP2 in the LD proteomes of the *sldp* mutant lines were assessed. As shown in Figure 3A, SLDP1 protein was nearly absent from LD-enriched fractions derived from *sldp1-1* and *sldp1-1 sldp2-1* mutant seedlings. Similarly, SLDP2 was not detected in the *sldp2-1* and *sldp1-1 sldp2-1* LD proteomes, but also in WT and *sldp1-1* derived LD proteomes it was only detectable in one of three replicates each. This is consistent with SLDP2 (and SLDP1) being a low abundant LD protein (Kretzschmar *et al*., 2020).

**Figure 3.**
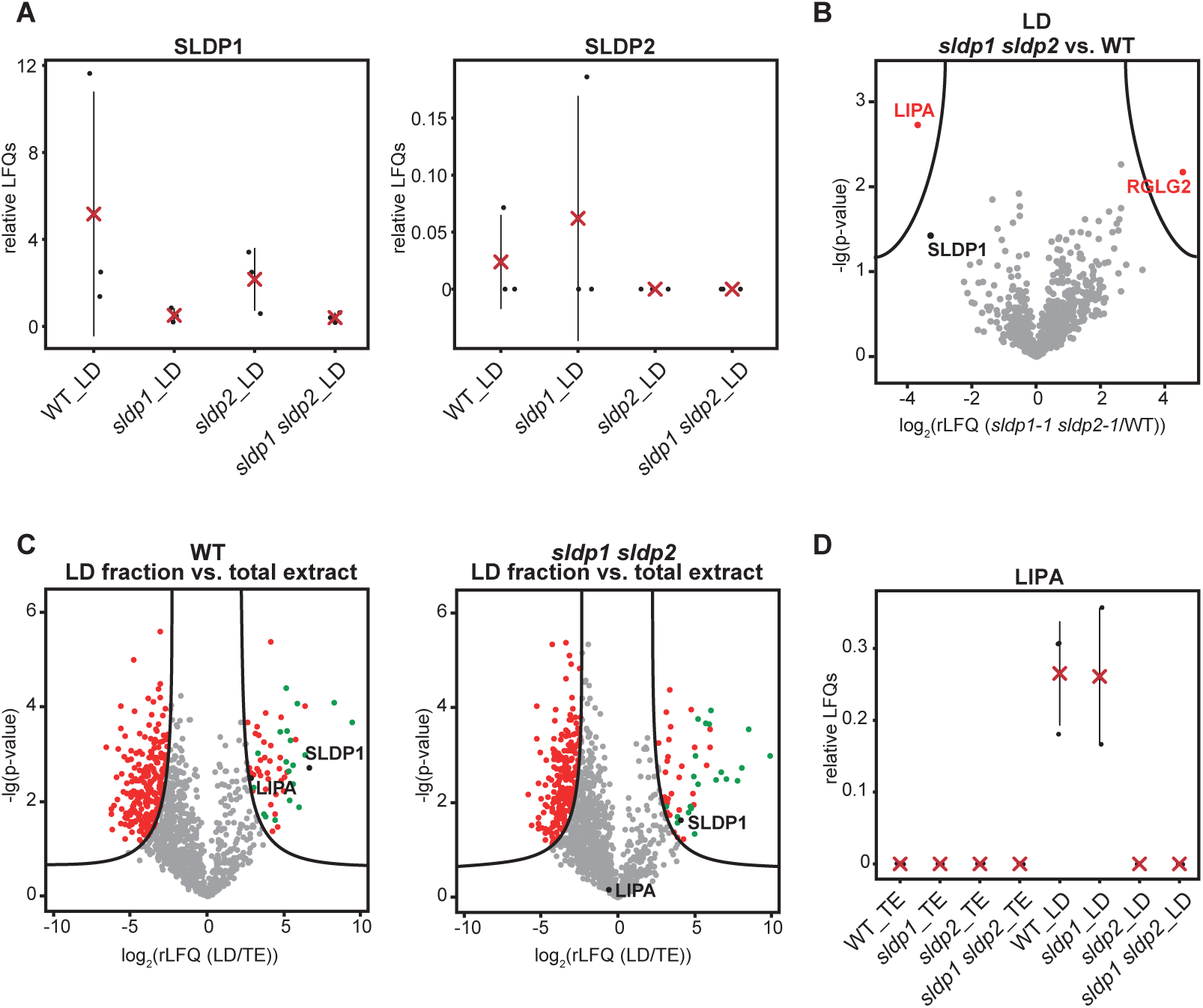
Proteomic analysis of *sldp* mutants. Proteins were isolated from germinating seedlings 36 h after stratification. Proteins from LD-enriched fractions and total protein fractions were analysed by LC-MS/MS after a tryptic in-gel digest (n=3 biological replicates). **A** rLFQ values of SLDP1 and SLDP2 analysed in LD-enriched fractions of different Arabidopsis lines (red cross = mean, lines = SD, n=3). **B** Volcano plot of imputed rLFQ values from LD fractions of the wild type versus *sldp1-1 sldp2-1*. It displays proteins differentially accumulating in the double mutant. Top left-hand corner proteins are significantly reduced in *sldp1-1 sldp2-1* LD fractions, top right-hand corner proteins are significantly reduced in wild-type LD fractions. **C** Volcano plots of imputed rLFQ values from TE versus LD fractions of wild-type and *sldp1-1 sldp2-1* seedlings, to detect proteins enriched at LDs. Proteins in the top right-hand corner are significantly enriched at LDs in the respective analysed line. SLDP1 and LIPA are marked in black and known LD-proteins among the significantly LD-enriched proteins are marked in green. **D** rLFQ values of LIPA analysed in LD- and TE-enriched fractions of different Arabidopsis lines (red cross, mean; lines, SD; dots, individual data points; n=3).

We then compared the TE and LD-enriched fractions to assess any differences in protein composition between WT and *sldp* mutant seedlings. For this, stringent filters (i.e., proteins only detected at least three times in at least one group and identified by at least two peptides) were applied to the whole data set (see Supplemental Data S2 for raw LFQs and S3 for normalised and filtered LFQs). Imputations were performed so that fold changes could also be calculated for proteins absent in one of the fractions (Supplemental Data S4). Comparisons of the TE fractions did not reveal any statistically significant differences between the WT and any of the *sldp* mutants (Supplemental Figure 10D). Likewise, comparisons of known LD proteins from LD-enriched fractions of WT and *sldp* mutants revealed no significant changes (Supplemental Data S4). However, two other proteins were found to be significantly differentially abundant in LD fractions of WT and *sldp* mutant seedlings: RING DOMAIN LIGASE2 (RGLG2) and LIPA (Figure 3B), with LIPA being absent in LD fractions from both *sldp2-1* and *sldp1-1 sldp2-1* seedlings and RGLG2 being more abundant in *sldp1-1 sldp2-1* seedling LD fractions (Figure 3B; Supplemental Figure 10E). Moreover, in WT and *sldp1-1*, LIPA was significantly enriched in LD fractions (i.e. absent from TE fractions, Figure 3C, D; Supplemental Figure 10E). As expected, a similar enrichment of LIPA was not found in LD fractions from *sldp2-1* and *sldp1-1 sldp2-1,* as LIPA was not detected in these samples (Figure 3D). Taken together, these data suggest that LIPA is associated with isolated LDs. They further indicate that SLDPs might act as a link between isolated LDs and LIPA and that LIPA is therefore important to mediate some of the functions of SLDPs. The link of SLDP to a higher abundance of RGLG2 is however harder to interpret. Furthermore, LIPA, in comparison to RGLG2 (Cheng *et al*., 2012; Yu *et al*., 2021), has previously not been explored in any detail. In conclusion, we aimed to further elucidate the function of LIPA, while the role of RGLG2 could be studied in the future.

### Co-expressed LIPA and SLDP2 mutually influence their subcellular localisation

Based on information provided by The Arabidopsis Information Resource (TAIR) (Berardini *et al*., 2015), LIPA has no annotated functions. While LIPA homologues exist in other plant species, they are absent in non-plant species, indicating that LIPA, like SLDP, is a plant-specific protein. LIPA is a 144 amino-acid-long protein with no described protein domains/motifs or putative transmembrane domains, nor any other obvious physicochemical features consistent with an LD targeting signal (e.g., putative amphipathic helix and/or hydrophobic sequence, Supplemental Figure 11A-C).

To assess the subcellular localisation of LIPA in plant cells, we transiently expressed the protein as an mVenus-tagged fusion protein in *N. tabacum* pollen tubes. As shown in Figure 4A, C-terminal mVenus-tagged LIPA (LIPA-mVenus) localised to the cytosol, while its N-terminal mVenus-tagged counterpart, mVenus- LIPA, localised predominantly to the PM. Notably, in *Agrobacterium*-transformed *N. benthamiana* leaves, another well-established plant cell model system for studying protein localisation (Sparkes *et al*., 2006), no apparent differences in localisation were observed between N- or C-terminal-GFP-tagged LIPA. Both LIPA fusion proteins localised to the PM and also the cytosol, which are closely appressed in these cells due to presence of the large central vacuole (Supplemental Figure 12A).

**Figure 4.**
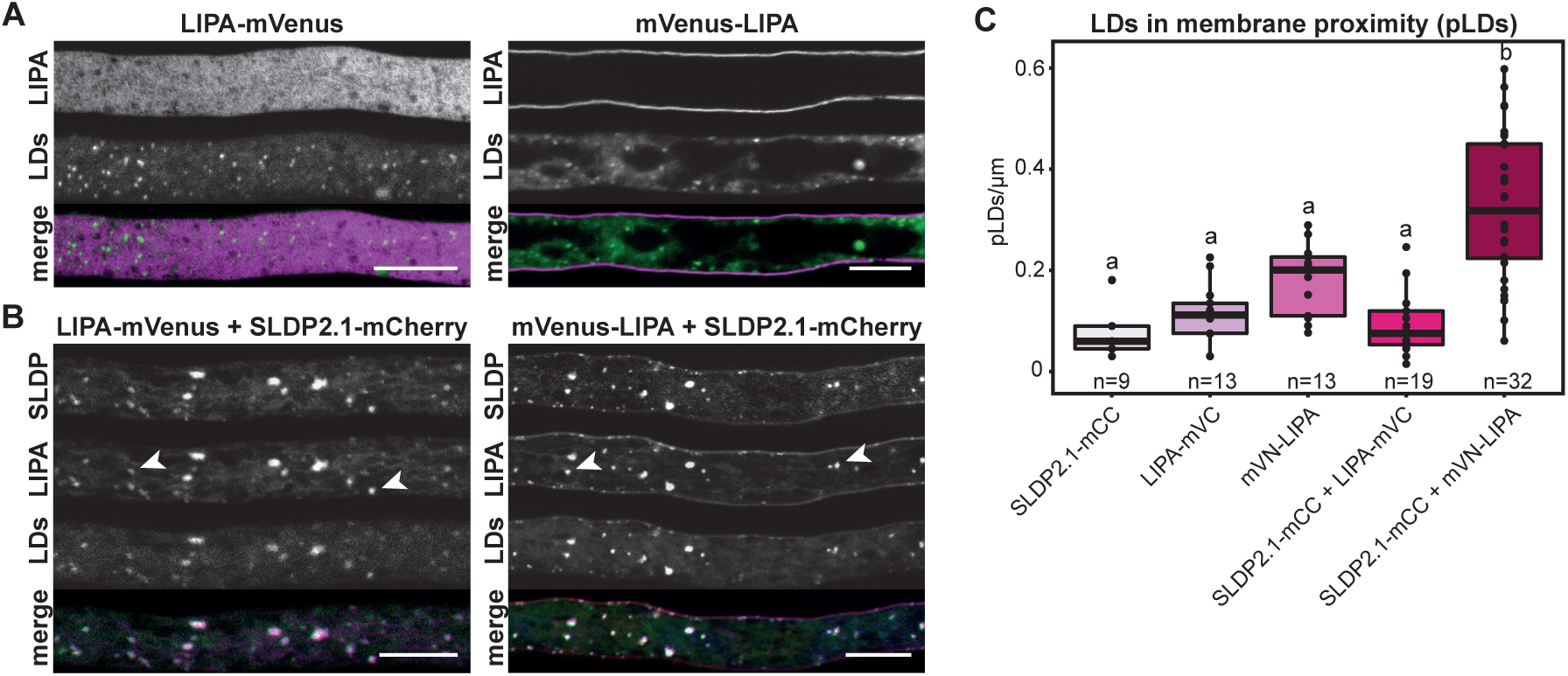
Subcellular localisation of LIPA in tobacco pollen tubes. **A, B** CLSM images of LIPA-mVenus and mVenus-LIPA transiently-expressed alone (**A**) or co-expressed (**B**) with SLDP2.1-mCherry in *N. tabacum* pollen tubes. LDs were stained with Lipi-Blue. LIPA colocalises with LDs only in the presence of SLDP2.1 (see arrowheads) but not when expressed alone. Images are representative of at least 5 micrographs of transformed pollen tubes with the indicated fusion construct(s). For merged images with two channels: magenta: mVenus (LIPA); green: LDs. For merged images with three channels: red: mVenus (LIPA), blue: mCherry (SLDP), green: LDs. Bars, 10 µm. **C** Analysis of LDs in proximity to the PM. Lipi-Blue-stained LDs adjacent to the PM (i.e., pLDs) in *N. tabacum* pollen tubes (refer to **A**, **B**) were counted manually and the number of pLDs per µm was calculated for each indicated construct(s). Results are presented as boxplots (displaying lower hinge = 25% quantile, median = 50% quantile, upper hinge = 75% quantile, upper whisker = largest observation less than or equal to upper hinge + 1.5 * IQR, lower whisker = smallest observation greater than or equal to lower hinge - 1.5 * IQR). One-way ANOVA was performed, followed by Tukey post-hoc analysis (F (4,81) = 23.37, p = 7.24e-13, n = as indicated). Note that only the SLDP2.1 + mVenus-LIPA co-bombardment increased the number of pLDs compared to the single bombardment controls. Statistical results are presented as compact letter display of all pair-wise comparisons.

While LIPA is localised to the PM (and cytosol) in transiently-transformed plant cells (Figure 4A, Supplemental Figure 12A), proteomic analysis indicated that LIPA co-purified with SLDP2-containing LDs, but not with LDs lacking SLDP2 (Figure 3B, Supplemental Figure 10E,F). Hence, LIPA appears to be a PM protein that also associates with LDs in an SLDP2-dependent manner. To further test this hypothesis, we co-expressed Arabidopsis SLDP2.1 and LIPA. *N. tabacum* pollen tubes, lacking homologues of both SLDPs and LIPA based on previous proteomic data (Kretzschmar *et al*., 2018), were chosen as expression system to avoid endogenous SLDP and/or LIPA from potentially interfering with localisation analyses. SLDP2.1 (and SLDP1.1) tagged with mCherry showed the same LD-localisation in pollen tubes as its mVenus-tagged counterparts (compare images in Supplemental Figure 13A and Figure 1C). As shown in Figure 4B, co-expression of SLDP2.1-mCherry and LIPA-mVenus (C-terminally tagged) in pollen tubes resulted in LIPA being re-located from the cytosol to LDs (compare with images of LIPA-mVenus expressed on its own; Figure 4A). A similar relocation was observed when LIPA-mVenus was co-expressed with either SLDP1.3-mCherry or the non-tagged (native) versions of SLDP1.3 or SLDP2.1 (Supplemental Figure 13B).

Next, we assessed the localisation of co-expressed SLDP2.1-mCherry and mVenus-LIPA (N-terminally tagged). As shown in Figure 4B, this co-expression resulted in a change of localisation of both proteins: co-expressed SLDP2.1 and LIPA were both dually localised to LDs and the PM (Figure 4B), unlike their localisation exclusively to LDs or the PM, respectively, when they were expressed on their own (Figure 4A, Figure 1B). Furthermore and as discussed below, a significant proportion of LDs decorated withSLDP2.1-mCherry and mVenus-LIPA appeared to be positioned close to the PM instead of distributed in the cytosol. By contrast, when mVenus-LIPA was co-expressed with SLDP1.3-mCherry, the proteins partially co-localised, but an increase in the association of LDs with the PM was not observed (Supplemental Figure 13B, Figure 4C). Likewise, in *N. benthamiana* leaves, SLDP2.1-mCherry localised to LDs when expressed on its own and both N- and C-terminal GFP-tagged versions of LIPA were re-located from the cytosol and PM to LDs when co-expressed with SLDP2.1-mCherry (Supplemental Figure 12B, C).

Taken together these data indicate that SLDPs and LIPA can influence each other’s localisation in plant cells, such that SLDP2.1 and SLDP1.3 can recruit LIPA to LDs, while LIPA can recruit at least SLDP2.1 and LDs to the PM.

### LIPA can immobilise SLDP2-containing LDs at the PM in pollen tubes

A consistent observation from experiments involving co-expressed SLDP2 and LIPA in pollen tubes (Figure 4B) was the distinct positioning of LDs at the PM in these cells. That is, in pollen tubes co-expressing SLDP2.1-mCherry and mVenus-LIPA, LDs appeared to be located in close proximity to the PM more often as compared to cells expressing either protein on its own (Figure 4B; compare also with images in Figure 1B and 4A). This suggests that SLDP2 and LIPA together are involved in the positioning of LDs, or at least a subset thereof, at the PM. To quantify this positioning, LDs in vicinity to the PM were manually counted, and put in relation to the pollen tube length in the taken micrographs for all transient expression combinations with SLDP2.1 and/or LIPA (Figure 4C). Consistent with the observations described above, co-expression of SLDP2.1-mCherry and mVenus- LIPA significantly increased the number of LDs in proximity to the PM, compared to either protein expressed alone. However, pollen tubes co-expressing SLDP2.1- mCherry and C-terminal-tagged LIPA-mVenus (or LIPA-mVenus alone) did not appear to differ in the number of LDs in proximity to PM, reinforcing our earlier conclusion that the C-terminus of LIPA is important for its association with the PM (Figure 4C).

To further test the premise that SLDPs and LIPA are important for the positioning of LDs at the PM, time-lapse imaging of Nile red-stained LDs in LIPA and SLDP2.1 co-transformed pollen tubes was performed. Pollen tube growth was extended by 3 h in comparison to previous experiments in order to give the tubes more time for protein expression and potential protein interactions. In control pollen tubes expressing mVenus alone (Supplemental Figure 13C, Supplemental movie S1), as well as in pollen tubes expressing mVenus-LIPA or SLDP2.1-mVenus alone (Figure 5, Supplemental movie S2-S3), LDs moved dynamically via cytoplasmic streaming. In contrast, in pollen tubes co-expressing mVenus-LIPA and SLDP2.1- mVenus, LDs were mostly localised and immobilised at the PM (Figure 5, Supplemental movie S4), as were, to a lesser extent, LDs in pollen tubes co-expressing SLDP1.3-mCherry and mVenus-LIPA (Supplemental Figure 13C, Supplemental movie S5-S6). Notably, the observed immobilisation of LDs at the PM was not due to the fluorophore appended to LIPA and SLDP. That is, co-expression of native (non-tagged) variants of LIPA and SLDP1.3 or SLDP2.1, along with mVenus alone serving as a cell transformation marker, yielded similar results and, in fact, even a more pronounced association of LDs with the PM in SLDP1.3 and LIPA co-expressing pollen tubes (Supplemental Figure 13C, Supplemental movies S7-S8).

**Figure 5.**
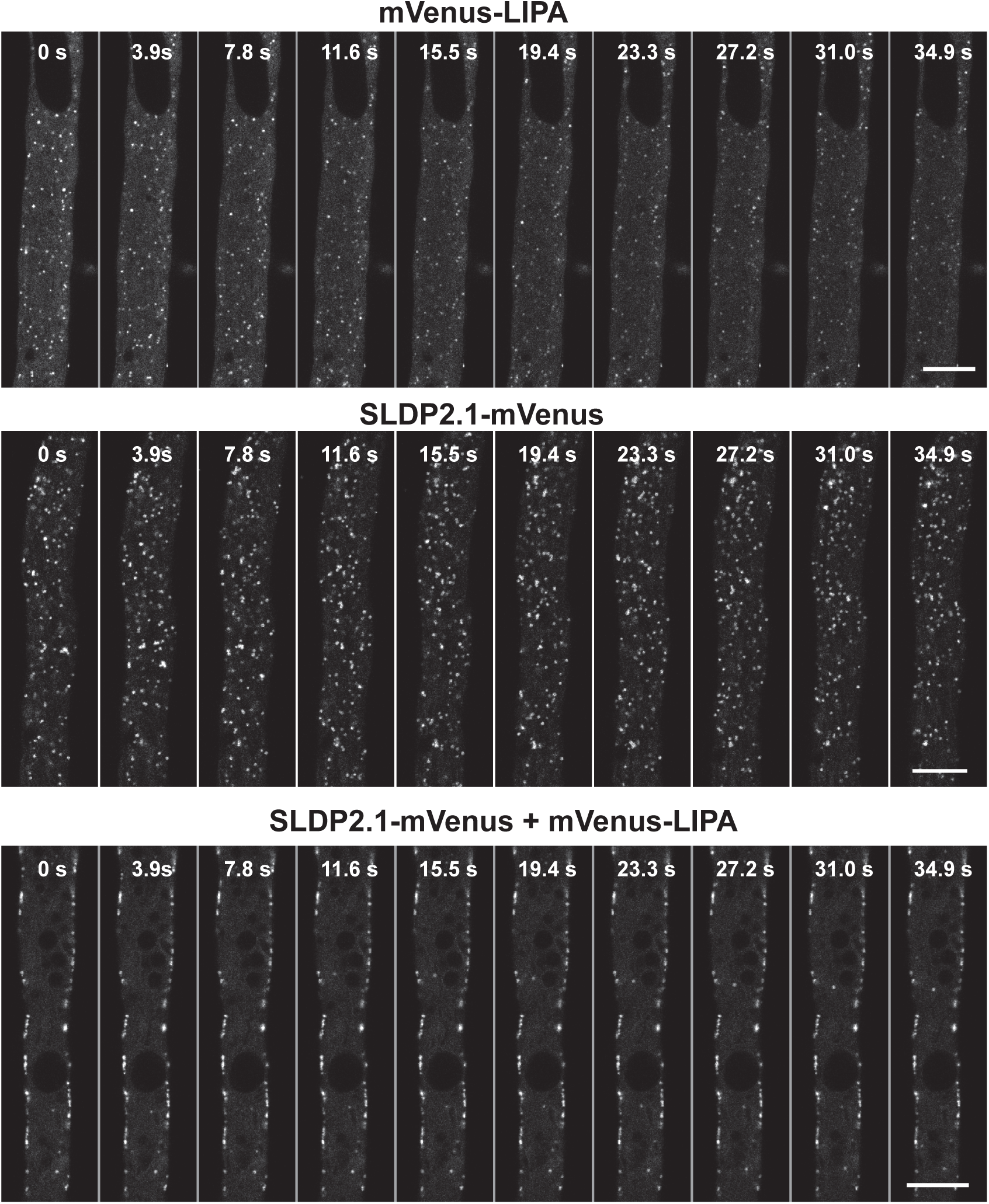
LD mobility analysis of LIPA and SLDP2.1 transformed tobacco pollen tubes. CLSM images of mVenus-LIPA and SLDP2.1-mVenus transiently expressed either alone or together in *N. tabacum* pollen tubes. Pollen tubes were stained with Nile red and LD dynamics were recorded over the indicated time course. Images are representative of time-course series of 5 transformed pollen tubes with each of the indicated fusion construct(s). Note in mVenus-LIPA and SLDP2.1-mVenus- transformed pollen tubes, LDs display dynamic cytoplasmic streaming, while in mVenus-LIPA and SLDP2.1-mVenus co-transformed pollen tubes LDs were predominantly immobilised at the PM. Bars, 10 µm.

### A coiled-coil domain in LIPA mediates its interaction with SLDP2 at LDs

While *in silico* analysis of Arabidopsis LIPA did not yield any known protein functional domains/motifs, the prediction program COILS (Lupas *et al*., 1991) revealed a putative coiled-coil domain in LIPA at residues 60–113 (Supplemental Figure 11D). Further structural predictions using the AlphaFold2 algorithm (Jumper *et al*., 2021; Varadi *et al*., 2022), which recently generated highly accurate structural models for eleven proteomes, including Arabidopsis, revealed that LIPA likely contains four α- helices (H1-H4), connected by flexible unstructured linkers (Figure 6A). H1, H2 and the majority of H3 are predicted with high confidence, while the shortest helix, H4, is of lower confidence and is a part of the otherwise unstructured C-terminus. The putative coiled-coil domain of LIPA corresponds to the helices H2 and H3.

**Figure 6.**
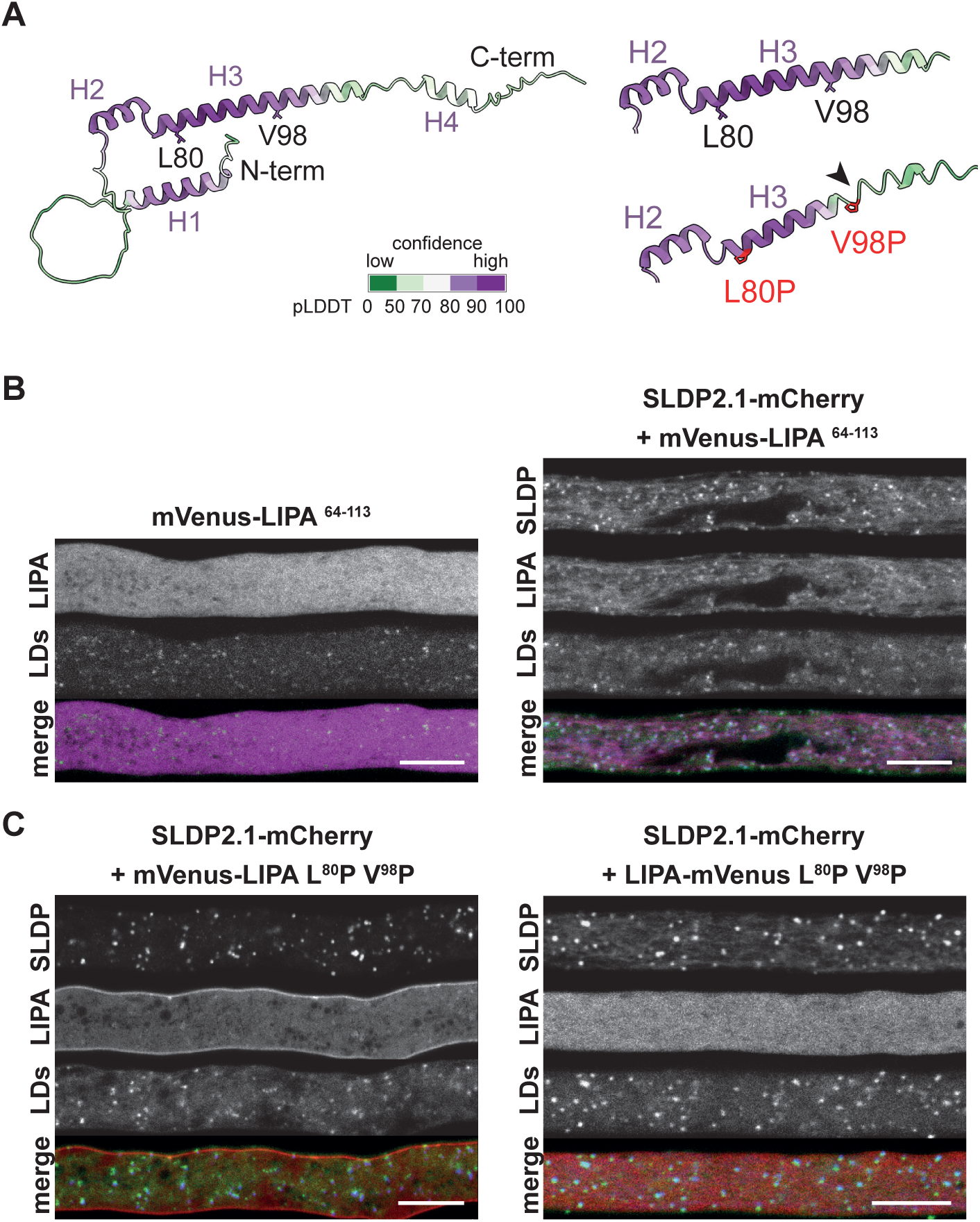
Analysis of the predicted α-helical/coiled-coil domain in LIPA. **A** Predicted structure of LIPA (left). The mutation of the L80 and V98 in helix 3 (right) to proline residues leads to a shortening of the helix (black arrow). Structures were generated using the AlphaFold2 algorithm. The models are coloured by local model confidence (pLDDT) as calculated by AlphaFold2. The pLDDT > 90 (dark purple) indicates regions of high prediction accuracy of both backbone and side chains. The pLDDT > 70 (white) indicates high-confidence backbone prediction. Regions in light and dark green (pLDDT < 70) represent low confidence predictions. **B**, **C** CLSM images of mVenus-LIPA^64-113^ and SLDP2.1-mCherry (**B**) and mVenus- LIPA L^80^P V^98^P or LIPA L^80^P V^98^P-mVenus and SLDP2.1-mCherry (**C**) transiently- expressed in *N. tabacum* pollen tubes. LDs were stained with Lipi-Blue. Images are representative of at least 5 micrographs of transformed pollen tubes with the indicated fusion construct(s). Note that mVenus-LIPA^64-113^, but not mVenus-LIPA- L^80^P V^98^P or LIPA-L^80^P V^98^P-mVenus re-locate to LDs upon SLDP2.1-mCherry co-expression. For merged images with two channels: magenta: mVenus (LIPA); green: LDs. For merged images with three channels red: mVenus (LIPA), blue: mCherry (SLDP), green: LDs. Bars, 10 µm.

To analyse the role of the putative coiled-coil domain in subcellular localisation of LIPA, a truncated version of LIPA, consisting of the coiled-coil domain alone appended to mVenus (i.e., mVenus-LIPA^64-113^), was expressed in pollen tubes, either on its own or together with SLDP2.1. As shown in Figure 6B, mVenus-LIPA^64-113^ expressed alone localised to the cytosol, but was recruited to LDs upon co-expression with SLDP2.1-mCherry, consistent with results of co-expressed full-length LIPA and SLDP2.1 (Figure 4A, B). This indicates that the coiled-coil domain of LIPA is sufficient for its SLDP-mediated relocation to LDs.

The importance of the putative coiled-coil region in LIPA was also assessed by replacing a leucine and a valine residue at positions 80 and 98, respectively, with prolines, in order to disrupt the putative coiled-coil structure (Chang *et al*., 1999; Cheng *et al*., 2001); refer to the COILS prediction of LIPA L^80^P V^98^P mutant protein shown in Supplemental Figure 11D). The predicted structure of LIPA L^80^P V^98^P mutant protein was also assessed using the AlphaFold2 algorithm (Mirdita *et al*., 2021, Preprint). Although the algorithm is generally not well suited for predicting effects of individual point mutations (Akdel *et al*., 2021, Preprint), helix H3 was still predicted to be significantly shorter in the mutant protein than in native LIPA (Figure 6A). Notably, LIPA L^80^P V^98^P mutant protein with either N- or C-terminally appended mVenus, irrespective of co-expression with SLDP, localised in a similar manner as its native LIPA counterparts, i.e., to the PM and/or cytosol, but not to LDs (Figure 6C, Supplemental Figure 14). Thus, the predicted coiled-coil region of LIPA appears to be both sufficient and necessary to induce LIPA-mediated relocation of SLDP2- containing LDs.

### Both FRET/FLIM and Y2H assays confirm SLDP-LIPA interaction

To further test the hypothesis that SLDPs and LIPA interact, both FRET-FLIM and Y2H analyses were performed.

FRET-FLIM experiments were carried out in tobacco pollen tubes. As shown in Figure 7A, the co-expression of SLDP1.3-mVenus or SLDP2.1-mVenus with mCherry-LIPA led to a significant decrease in the fluorescence lifetime of mVenus in comparison to the expression of the SLDP1.3-mVenus and SLDP2.1-mVenus on their own. These results indicate that SLDP and LIPA come in close proximity at the surface of LDs. The putative interaction of SLDP and LIPA was also assessed testing truncated versions of SLDP1 and 2 that mislocalise to the cytosol in pollen tubes, i.e., SLDP1.3^Δ1-81^ and SLDP2.1^Δ1-75^ (refer to Figure 1B). Similar to full-length SLDP1.3- mVenus and SLDP2.1-mVenus, co-expression of SLDP1.3^Δ1-81^ or SLDP2.1^Δ1-75^ with LIPA-mCherry, led to a significant reduction in the fluorescence lifetime of mVenus, while co-expression with mCherry alone did not (Figure 7B). These results indicate that the N-termini of the SLDPs are not required for the interaction with LIPA and that an interaction does not depend on the localisation to LDs.

**Figure 7.**
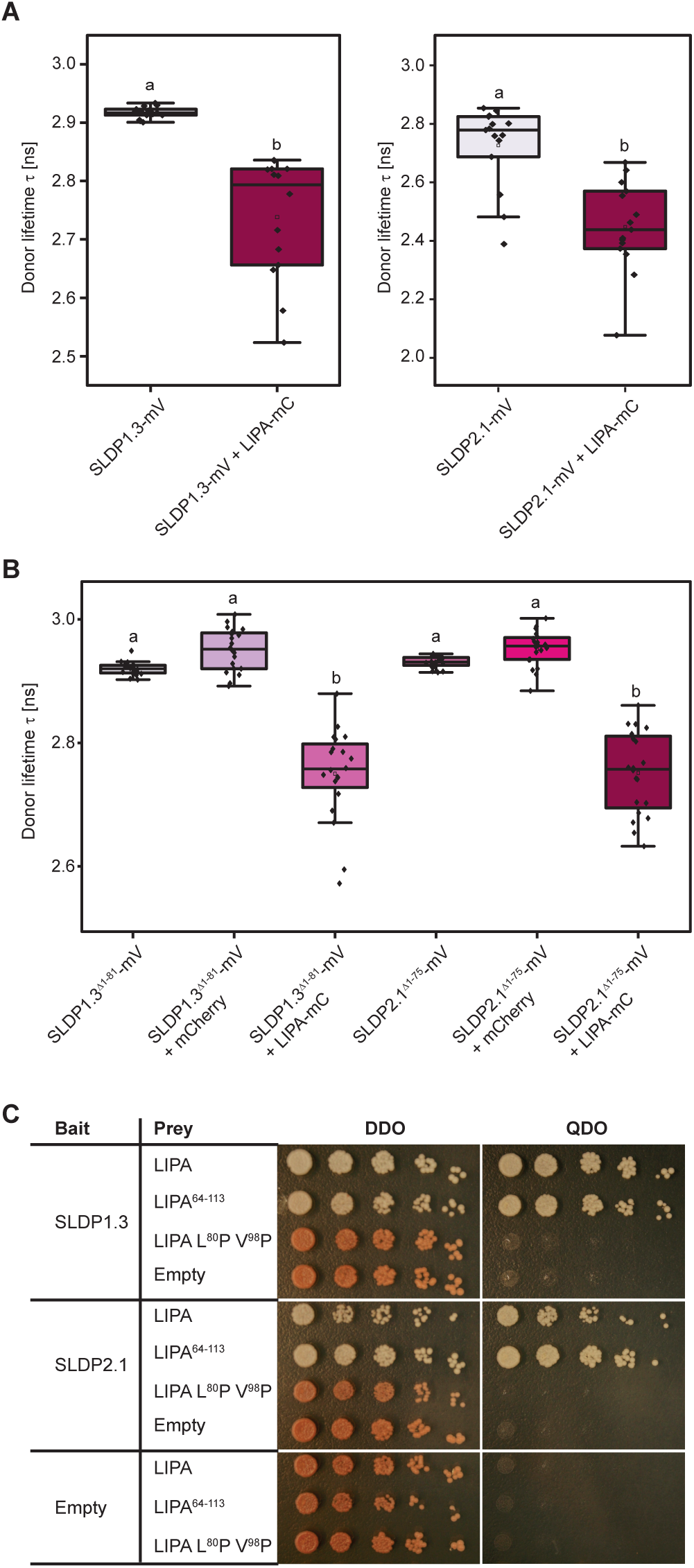
SLDP and LIPA interaction assays by FRET-FLIM and Y2H. **A** Full-length versions of SLDPs tagged with mVenus (mV) were expressed in tobacco pollen tubes either alone or in combination with the cytosolic LIPA-mCherry (LIPA-mC). Co-expression led to a recruitment of LIPA-mCherry to the LDs and a significant reduction of the donor lifetime. Fig7A: One-way ANOVA was performed, followed by Tukey post-hoc analysis (left panel: F (1,27) = 43.85, p = 4.18e-07, n1=15, n2=14; right panel: F (1,28) = 27.33, p = 1.49e-05, n = 15. Statistical results are presented as compact letter display of all pair-wise comparisons. **B** The expression of truncated cytosolic versions of the SLDPs with cytosolic LIPA- mCherry also led to a reduction of the donor lifetime in comparison to expression of the SLDPs alone, or of the SLDPs in combination with mCherry. One-way ANOVA was performed, followed by Tukey post-hoc analysis (F (5,114) = 94.57, p = 6.68e- 39, n=20). Statistical results are presented as compact letter display of all pair-wise comparisons. **C** Y2H interaction analysis of SLDP1, SLDP2 and LIPA. Yeast (*S. cerevisiae*) were co-transformed with bait (pGBKT7) plasmids containing full-length SLDP1 or SLDP2 and prey (pGADT7) plasmids containing LIPA or modified versions thereof, or with the corresponding empty plasmids serving as negative controls. Serial dilutions of transformed yeast cell cultures were then plated onto either plasmid-selection conditions (double drop out medium, DDO), or higher stringency selection conditions (quadruple drop out medium, QDO) where yeast cell growth requires a Y2H protein-protein interaction. Note that only yeast cells co-expressing SLDP1 or SLDP2 and LIPA or LIPA^64-113^, but not LIPA L^80^P V^98^P, grew on QDO plates. Results shown are representative of three separate co-transformations of yeast with each plasmid combination.

Additionally, the interaction of SLDP and LIPA or mutant versions thereof was also addressed in Y2H assays. As shown in Figure 7C, results showed that both, SLDP1.3 and SLDP2.1, interact with full-length LIPA or the putative coiled-coil domain of LIPA (LIPA^64-113^), but not with LIPA L^80^P V^98^P (Figure 7C). As expected, yeast expressing either SLDP1, SLDP2 or LIPA with only the corresponding ‘empty’ vector did not grow on selection media. Further, Y2H assays revealed that SLDP1.3, SLDP2.1 and LIPA do not self-associate, nor do SLDP1.3 and SLDP2.1 associate with each other (Supplemental Figure 15).

### LIPA targets the PM via its C-terminal polybasic region

Our experiments suggest that the putative coiled-coil region of LIPA (residues 60-113) is involved in binding SLDPs, but is not sufficient for its localisation to the PM (Figure 6B). In order to determine the region(s) in LIPA required for PM targeting, several truncated versions of the protein were generated and expressed as N-terminal mVenus fusions in tobacco pollen tubes. As shown in Figure 8A, the C-terminus of LIPA (residues 107-144), which includes a polybasic region (residues 107-134; Figure 8B), localised to the PM similar to full-length LIPA. By contrast, a shorter C-terminal region of LIPA (residues 120-144) was mislocalised to the cytosol, indicating that the polybasic region in LIPA is necessary for its PM targeting (Figure 8B). We also removed additional amino acids from the C-terminal end of the LIPA in the context of the LIPA^107-144^ mutant, including the C-terminal cysteine residue (LIPA^107-143^), which could potentially serve as a lipid-anchor site, and the C-terminal 10 or 21 amino acids (LIPA^107-134^ and LIPA^107-123^, respectively). Overall, the localization results for these LIPA mutants indicated that the C-terminal 10 amino acids are not essential for PM targeting, while deletions of residues within the polybasic region abolished the PM targeting of LIPA (Figure 8B). Taken together, the amino acids 107-134 are able to bind the PM.

**Figure 8.**
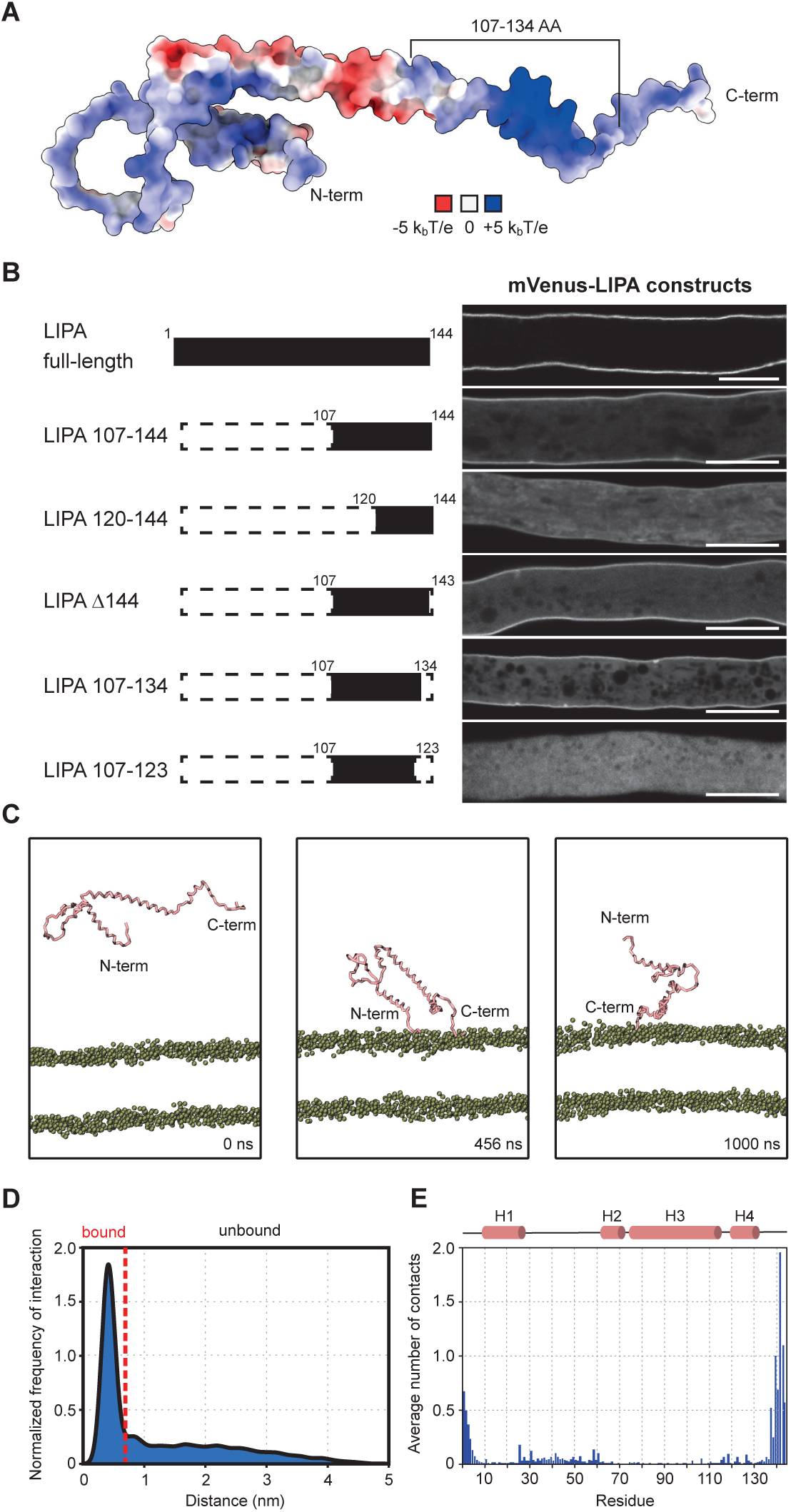
PM-localisation of LIPA. **A** Electrostatic potential mapped onto the solvent-excluded surface of the LIPA structure. Charge distribution indicates a strong accumulation of positively charged residues in the C-terminal region, especially region 107-134. **B** Illustrations and CLSM images of full-length and various truncation versions of LIPA appended to mVenus in transiently-transformed *N. tabacum* pollen tubes. Images are representative micrographs of at least 10 transformed pollen tubes. Bars, 10 µm. **C** Snapshots from the molecular dynamics (MD) simulations. Different time points are indicated. The protein is shown in the ribbon representation (pink). Only phosphate groups of the lipid bilayer are shown for the sake of clarity. **D** Probability density distribution of protein-membrane minimum distances shows a significant portion of the bound protein to the lipid bilayer. **E** Mean number of contacts between protein and phosphate group of the lipid bilayer. The contacts were defined as the number of phosphate groups within 0.8 nm of protein atoms. The C-terminus displays the highest number of contacts, but helix H1 and the adjacent linker also contribute to the interaction. The secondary structure is indicated above the plot.

To further investigate the interaction of LIPA with the PM at the molecular level, we utilised coarse-grained molecular dynamics (MD) simulations. This computational approach has been successfully used to study the interaction of a wide variety of membrane proteins with a lipid bilayer (Corradi *et al*., 2019; Marrink *et al*., 2019). Here, we used the recently released version (3.0) of the Martini force field (Souza *et al*., 2021), which was shown to accurately describe the membrane-binding behaviour of several membrane proteins and to correctly identify pivotal amino acid residues involved in the interaction (Srinivasan *et al*., 2021). To study LIPA, the simulated system contained one molecule of LIPA, ions, water molecules, and a complex phospholipid bilayer with a composition mimicking the negatively-charged plant cell PM (Wassenaar *et al*., 2015). We performed five independent MD simulations with different starting velocities, resulting in a total of 5 μs simulation time. During the simulations, LIPA displayed an on/off membrane binding similarly to the behaviour reported for other membrane proteins (Srinivasan *et al*., 2021) (Supplemental Figure 16). Figure 8C shows selected snapshots from the MD simulations depicting unbound and membrane-bound states of LIPA (See also Supplemental movie S9). We then quantified the binding events by generating a probability density distribution using the kernel density estimation method and, as shown in Figure 8D, the calculated probability density distribution revealed a significant population of LIPA in the membrane-bound state. Next, we investigated amino acid residues of LIPA potentially involved in the interaction with negatively charged phospholipids. In agreement with the results from our truncation analyses of LIPA (Figure 8A), the highest number of contacts is located at the C-terminus of LIPA (Figure 8E). In addition, we also observed a contribution of H1 and, to a lesser extent, of the adjacent flexible region, to the interaction of LIPA with negatively charged phospholipids (Figure 8E). Interestingly, the region corresponding to the helical/coiled-coil domain (i.e., H3) involved in the LIPA-SLDP interaction (Figure 6), showed no interaction with the phospholipids (Figure 8E), further corroborating the role of this region in protein-protein interactions (Figure 8E).

### *lipa* mutants phenocopy *sldp1 sldp2* mutants in their aberrant subcellular distribution of LDs during post-germinative seedling growth

Given that SLDP and LIPA appear to act together in the positioning of LDs at the PM, we next tested whether disruption of *LIPA* would have a similar effect on the subcellular distribution of LDs, as observed upon disruption of *SLDP* (Figure 2). To this end, two independent Arabidopsis mutant lines of *LIPA* were generated, a T-DNA insertional line, *lipa-1*, and a CRISPR/Cas9 deletion line, *lipa-2,* which is devoid of most of the *LIPA* open reading frame (Figure 6A, Supplemental Data S1). RT-qPCR analyses confirmed a lack of full-length *LIPA* transcripts in both mutant lines (Supplemental Figure 17). Time-course imaging of Nile red-stained LDs in cotyledon and hypocotyl cells in WT and both *lipa-1 and lipa-2* rehydrated (mature) seeds and seedlings after stratification, was performed as described above for the *sldp* mutants (Figure 9B, Supplemental Figure 6; compare with images presented in Figure 2B). While again no obvious differences in LD distribution or storage vacuole appearance were observed in WT and *lipa* mutant rehydrated seeds (Supplemental Figure 7), an LD-clustering phenotype similar to *sldp2* and *sldp1 sldp2* seedlings at 24 and 36 h was readily observed (Figure 9B, Supplemental Figure 6B). Additionally, as described for *sldp* mutants, Z-stack images of cotyledon and hypocotyl cells in *lipa* mutants at the 36-h time point and quantification of the LD distribution in these cells were performed and revealed significant differences in the LD distribution as compared to WT cells (Supplemental Figure 8). These results provide further evidence that SLDP2 and LIPA act together in the proper positioning of LDs at the PM during post-germinative seedling growth.

**Figure 9:**
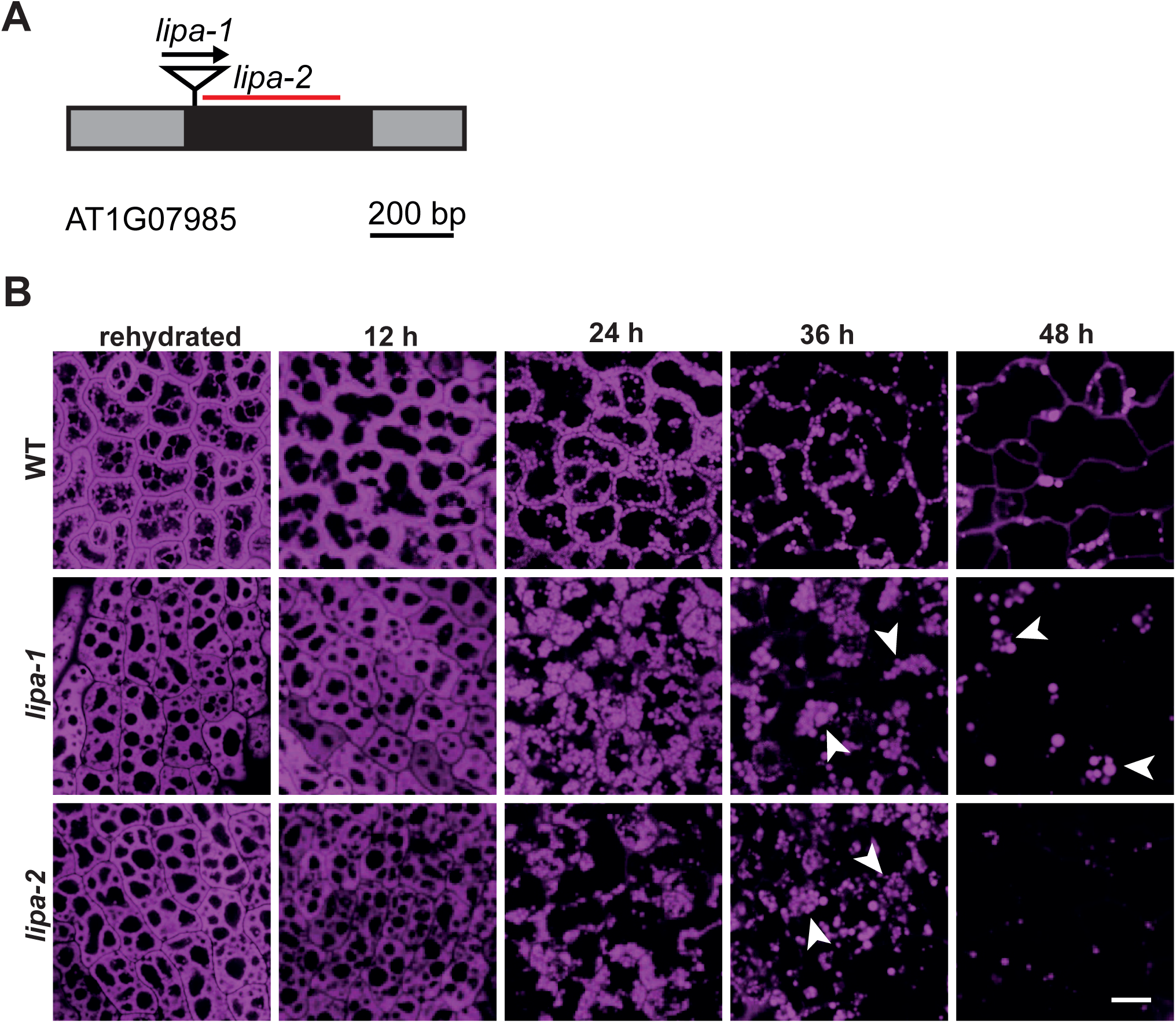
Time-course analysis of LDs in cotyledons. **A** Schematic depiction of the LIPA gene with untranslated regions (grey boxes), one exon (black box), T-DNA insertion site (triangle, arrow indicating direction of T-DNA) and the region deleted by CRISPR/Cas9 genome editing (red line). **B** CLSM images of rehydrated seeds and seedling cotyledon cells from WT and *lipa-1* and *lipa-2* mutant Arabidopsis lines. Seeds were rehydrated for 1 h or stratified for 4 days at 4 °C in the dark. LDs were stained with Nile red after rehydration, or 12, 24, 36 and 48 h (± 2 h) after stratification. Arrowheads indicate obvious examples of LD clusters in *lipa* mutant seedling. Images are single plane images from the middle of the cell (similar planes were chosen for all images). Images are representative of at least five micrographs of seeds and seedlings for each plant line and time point. Bar, 10 µm and applies to all images in the panels.

### Both mGFP-LIPA and SLDP-mCherry localise to contact sites between LDs and the PM

Previous results hint at a putative MCS between LDs and the PM, formed through interaction of LIPA and SLDP. On that account, we next investigated whether these proteins are specifically enriched at LD-PM contact sites in seedlings. Therefore, Arabidopsis transgenic lines stably expressing *SLDP1.3* and *SLDP2.1* (under control of the 35S promoter) appended to a C-terminal mCherry were assayed for their subcellular localisation in hypocotyl cells of 40-h old seedlings. As shown in Figure 10A and 10B, both SLDP1.3-mCherry and SLDP2.1-mCherry localised to the surface of LDs, as evidenced by the torus-shaped fluorescence patterns surrounding the fluorescence attributable to the BODIPY-stained neutral lipids inside the LDs. However, SLDP fluoresence, particularly SLDP2.1-mCherry, was often enriched at distinct sites on the LD surface that were presumably adjacent to the PM (Figure 10 B), suggesting that espeially SLDP2.1 preferentially localises at LD-PM contact sites. Similarly, eGFP-LIPA stably expressed in the Arabidopsis *lipa-1* background localised to the surface of LDs in hypocotyl cells and was often enriched at apparent LD-PM contact sites (as shown by staining of LDs with Nile Red and the PM with FM4-64).

**Figure 10.**
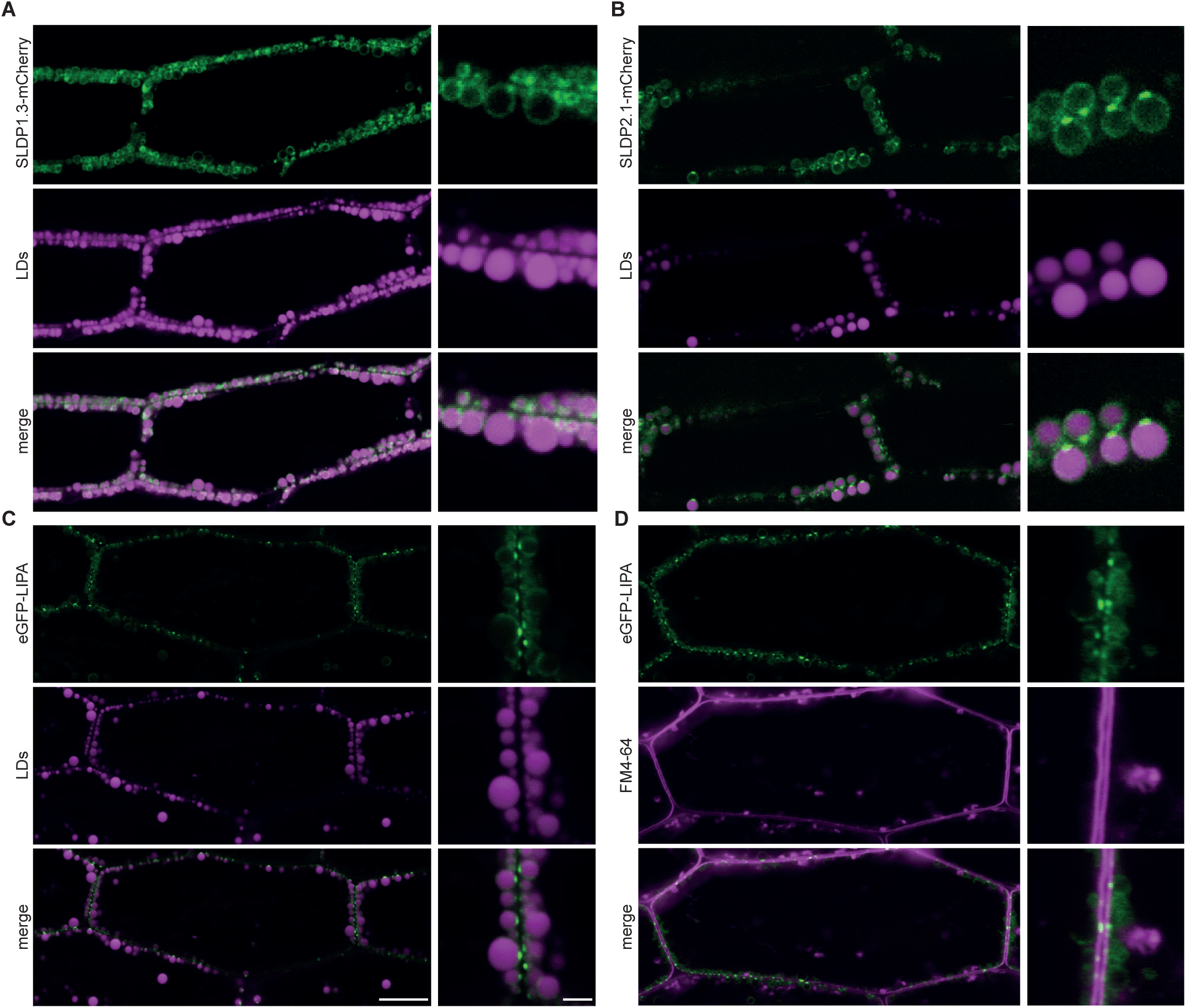
Localisation of stably-expressed SLDP and LIPA in Arabidopsis seedling hypocotyls. SLDP1.3-mCherry and SLDP2.1-mCherry (**A**, **B**) or eGFP-LIPA (**C**, **D**) were stably-expressed under the 35 S promoter in Arabidopsis Col-0 (**A**, **B**) or *lipa*-1 mutant (**C**, **D**) plants. Fusion protein localisation was monitored in 38 h-old seedlings by CLSM after staining with either BODIPY 493/503 (**A**, **B**), Nile Red (**C**), or FM4-64 (**D**). The panels on the right display portions of the cells at higher magnification in the panels to the right. Note that the fluorescence attributable to SLDP1.3-mCherry predominantly encircled LDs (**A**), while for SLDP2.1-mCherry and eGFP-LIPA, fluorescence was enriched at putative LD-PM MCSs (**B**-**D**). Images are representative of at least five seedlings from each of three (**A**, **B**) or two independent plant lines (**C**, **D**). Bars, 10 µm and 2 µm in low and high magnified images, respectively, and applies to all the corresponding images in the other panels.

## DISCUSSION

While plant LD research has yielded a number of significant advancements in recent years (Lundquist *et al*., 2020; Ischebeck *et al*., 2020; Kang *et al*., 2021), many important questions related to plant LD biology remain unanswered, including if and how they interact with other organelles and subcellular structures. Here, we took advantage of the recent proteomics-based identification of several novel LD-associated proteins (Kretzschmar *et al*., 2020) and characterised two members of the plant-specific SLDP family, SLDP1 and SLDP2. We showed that the LD-association of SLDPs is mediated by an N-terminal predicted amphipathic and hydrophobic region (Figure 1), similar to LD targeting sequences reported previously for other LD-localised proteins (Wilfling *et al*., 2013; Kretzschmar *et al*., 2018; Pyc *et al*., 2017). Moreover, in Arabidopsis *sldp* mutant seedlings, LDs display an aberrant subcellular distribution, with LDs clustering in the centre of the cell and not, as observed in wild-type seedlings, aligning along the PM (Figure 2).

We further showed that LIPA is associated with isolated LDs, depending on the presence of SLDP2 (Figure 3) and LDs in Arabidopsis *lipa* mutant seedlings displayed an aberrant clustering phenotype, similar to that observed in *sldp* mutant seedlings. Microscopic analyses revealed that transiently expressed LIPA localises to the PM in pollen tubes, but, upon co-expression with SLDP, is also found at LDs. Conversely, SLDP is partially re-located to the PM upon co-expression with LIPA (Figure 4). Moreover, we observed that at least a subset of LDs in LIPA and SLDP co-expressing pollen tubes are conspicuously immobilised at the PM and not streaming throughout the cytoplasm (Figure 4). How exactly LIPA associates with the PM remains unclear, as no putative transmembrane domains were detected within the LIPA protein sequence. Based on structural modelling and truncation analyses, though, we suggest that LIPA might bind the plasma membrane through electrostatic interactions of a positively charged polybasic sequence near LIPA’s C-terminus (in the region of amino acids 107-134) which could interact with negatively charged head groups of anionic lipids (Noack and Jaillais, 2020). We cannot rule out, however, that PM-association of LIPA might also require interaction with additional PM-localised protein(s), or that these might additionally stabilise the interaction (given our modellings display LIPA as a flexible protein that dynamically binds the PM (Figure 8C).

Lastly, we found that SLDPs and LIPA were enriched at contact sites between LDs and the PM in seedlings (Figure 10). Based on these and other results, we propose a working model (Figure 11) in which LIPA associates with the PM via a C-terminal region and, based on Y2H and FRET/FLIM experiments (Figure 7), likely directly interacts with SLDPs through its putative coiled-coil domain, a protein structural domain well-known to be involved in mediating protein-protein interactions (Mier *et al*., 2017). SLDPs in turn are anchored to LDs via their N-termini and then, through interaction with LIPA, tether LDs to the PM (i.e., forming an LD-PM MCS) during post-germinative seedling growth.

**Figure 11.**
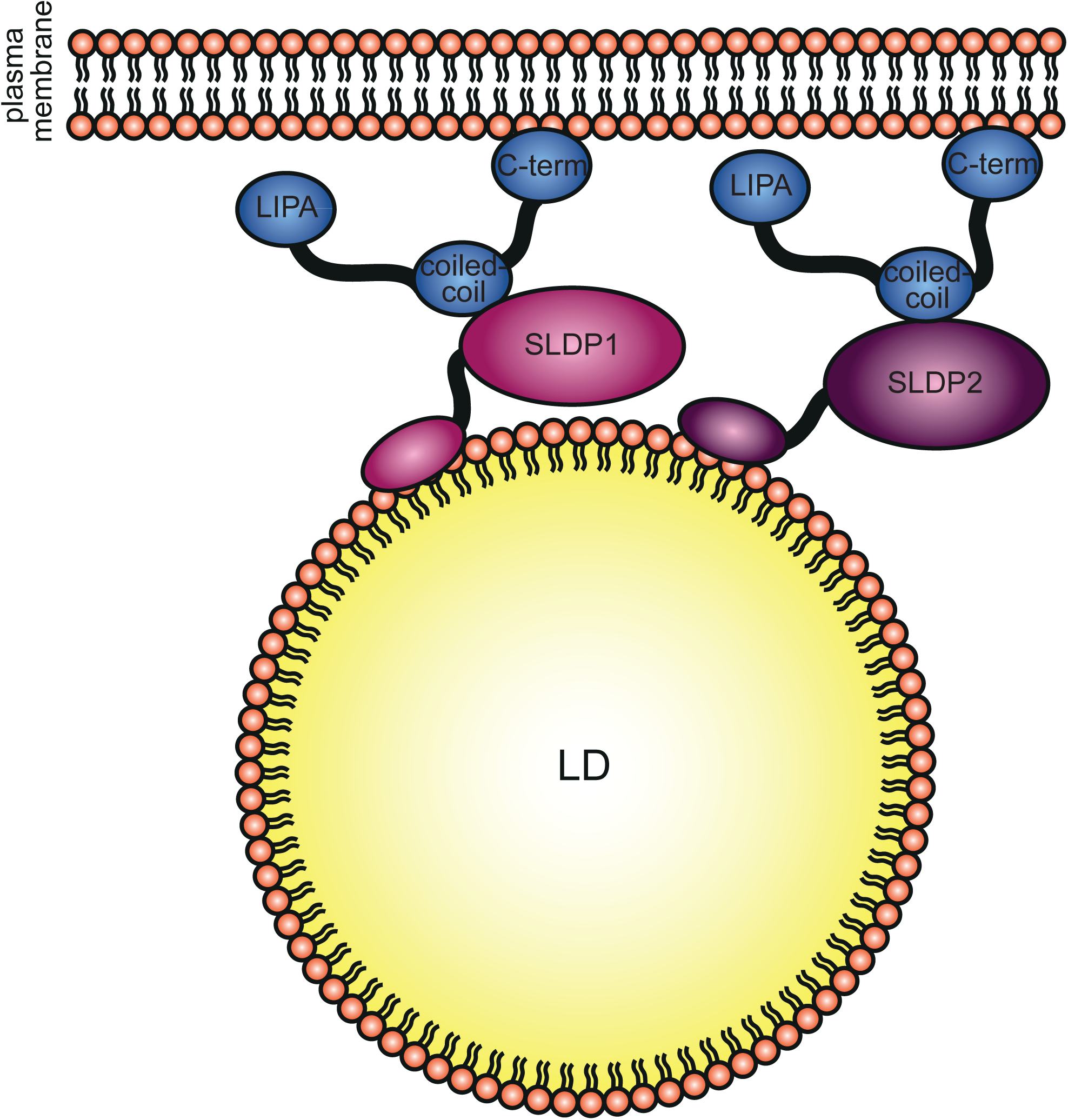
Model of SLDP-LIPA mediated PM-LD MCS. SLDP1 and SLDP2 associate with the surface of LDs via their N-terminal regions. LIPA binds to the PM through a C-terminal region and interacts with SLDP1 and SLDP2 via its coiled-coil region. The resulting SLDP1/2-LIPA interaction(s) tether the LD to the PM.

As to the function of a putative LD-MCS formed through interaction of SLDP and LIPA, we can only speculate. LDs are lipid storage sites and MCS of LDs are often involved in lipid transfer processes (Salo *et al*., 2019; Bohnert, 2020). In this regard, LDs in plant seedlings have mostly been considered as a source of acyl chains to fuel seedling establishment (Yang and Benning, 2018). Consequently, previous studies on LD-organelle MCS in plants have focused primarily on how LD-peroxisome MCS help facilitate the mobilisation of stored lipid reserves in LDs via peroxisomal β-oxidation (Esnay *et al*., 2020). Our results extend this work by showcasing a possible additional LD-involving MCS during post-germinative seedling growth: LDs in seedlings might be needed at the PM, either to provide lipids or to buffer (i.e., store) excess and potentially cytotoxic lipids produced during membrane repair and/or growth. More generally, LD-PM MCS might be required for maintaining PM lipid homeostasis, as has been shown for e.g. LD-ER MCS (Velázquez *et al*., 2016). The importance of this LD-PM MCS might therefore only come into effect upon stress conditions (such as salt, freezing, mechanical, etc.), when membrane composition has to be remodelled. This would explain the lack of any obvious growth and/or developmental phenotypes in the *sldp* and *lipa* mutants, which were examined under laboratory conditions in this study, despite their striking cellular (LD) phenotype.

Recently, several tri-organellar contact sites involving LDs and the ER have been described in mammals, yeast and insects (Freyre *et al*., 2019; Hariri *et al*., 2019; Ugrankar *et al*., 2019). As LDs are often associated with the ER (Hugenroth and Bohnert, 2020), another interesting question arising is whether the ER might be involved in the observed LD-PM MCS, as well. While the work presented here does not provide direct evidence for this hypothesis, it was recently found that SEIPIN, an ER-membrane protein that participates in LD biogenesis (Sui *et al*., 2018), interacts with the ER-associated protein VAP27-1 (vesicle-associated membrane protein (VAMP)-associated protein 27-1) at ER-LD MCS (Greer *et al*., 2020). VAP27-1 is also involved in tethering the ER to the PM via interaction with SYT1 (Siao *et al*., 2016), a homologue of mammalian extended synaptotagmins, which is known to be important during abiotic stress responses (Yamazaki *et al*., 2008; Schapire *et al*., 2008). Given that mammalian extended synaptotagmins are involved in lipid transfer (Schauder *et al*., 2014; Yu *et al*., 2016), a three-way MCS involving LD, ER and PM could serve as a key hub for lipid homeostasis at the plant PM.

Future work will now be aimed at investigating these possibilities by uncovering the mechanistic details underlying the putative SLDP-LIPA tethering complex reported here, as well as elucidating the physiological importance of LD-PM MCSs in seedlings and other tissues.

### Experimental Procedures

#### Plant material and growth conditions

All *Arabidopsis thaliana* (L.) plants employed the ecotype Col-0 or were derived from it in the case of T-DNA and CRISPR/Cas9 mutant lines. They were grown in a climate chamber (York) in 60 % relative humidity, with a constant temperature of 23 °C and under a 16 h/8 h day/night cycle with a daytime light intensity of 150 µmol photons m^−2^ s^−1^ (the climate chamber was equipped with LuxLine Plus F36W 830 Warm White de Luxe fluorescent tubes; Osram Silvania). Plants were either grown on soil or on half-strength MS medium (Murashige and Skoog, 1962) supplemented with 0.8 % (w/v) agar with or without 1 % (w/v) sucrose (as indicated) and stratified for four days at 4 °C in the dark. Seeds grown on medium were surface sterilised in 6% sodium hypochlorite solution for 15-20 minutes. For hygromycin selection, half-strength MS plates were supplemented with 25 µg/ml hygromycin and 1 % (w/v) sucrose, stratified for two days at 4 °C, subjected to light for 4 h and then kept vertically in the dark for three days. Hygromycin-resistant seedlings were transferred to half-strength MS + 1 % (w/v) sucrose without hygromycin for one week prior to transplanting them into soil.

Tobacco (*Nicotiana tabacum* L. cv. Samsun-NN) plants were grown in the greenhouse as previously described in order to collect pollen (Rotsch *et al*., 2017). Plants were kept under 14 h of light from mercury-vapor lamps in addition to sunlight. Light intensities reached 150 – 300 µmol m^−2^ sec^−1^ at the flowers and 50 – 100 µmol m^−2^ sec^−1^ at leaves at mid-height. Temperature was set to 16 °C at night and 21 °C during the day with a humidity of 57–68%.

*Nicotiana benthamiana* plants were grown in soil at 22 °C with a 16-h/8-h day/night cycle and 50 µE·m^-2^s^-1^ light intensity.

#### T-DNA lines

Knockout lines of *SLDP1, SLDP2* and *LIPA* were generated. The commercially available T-DNA insertional lines SALK_204434C (*sldp1-1*, T-DNA inserted in intronic region behind base 1028) and SALK_068917 (*sldp2-*1, T-DNA inserted in first exon behind base 42) and Gabi-KAT 723C08 (*lipa-1*, T-DNA inserted behind base 20) were used, and CRISPR/Cas9 was used to generate *sldp1-2*, *sldp2-2, and lipa-2 (see below)*. Sequence alignments and predicted protein products of all analysed mutant lines are shown in Supplemental Data S1.

#### CRISPR/Cas9

To generate CRISPR/Cas9 mutants, sgRNAs were designed using the Cas-Designer and Cas-OFFinder at http://www.rgenome.net/ for a SpCas9 protospacer adjacent motif (PAM) sequence and with a length of 19 bp (without PAM) against the *Arabidopsis thaliana* (TAIR10) genome (Bae *et al*., 2014; Park *et al*., 2015). Cloning was performed as described previously (Xing *et al*., 2014). As template for the sgRNA cassette (including one sgRNA backbone, one U6-26 terminator and one U6- 29 promoter), pCBC DT1T2 was used and the generated PCR-product was cloned into pHEE401E via BsaI restriction sites, between a U6-26 promoter on one side and a second sgRNA backbone and a U6-29 terminator on the other side (as described previously (Xing *et al*., 2014; Wang *et al*., 2015). This way, a CRISPR/Cas9 construct containing two sgRNAs under two U6 promoters and a Cas9 under the egg-cell specific EC1.2 promoter was obtained. To knock out one gene, two different sgRNAs were targeted at it, aiming at deleting the whole gene stretch between the target sequences. This made it possible to screen for mutant plants via PCR. For this, gDNA was extracted from rosette leaves, the area of interest was amplified via REDTaq®-PCR and screened for the desired smaller PCR-products that indicated a deletion. Homozygous mutants were obtained in the T2 and T1 generation for SLDP1 and SLDP2, respectively. To remove the Cas9-transgene, homozygous mutants were backcrossed to WT plants (and Cas9-loss was confirmed by PCR with U6- and Hygromycin resistance gene-specific primers).

For *SLDP1*, a mutant line with deletion of bases 333-564 in the first exon (resulting in a frameshift and a premature stop codon at position 650-652 for AT1G65090.1 and .2 or at position 686-688 for AT1G65090.3, producing a potential 139 amino acid protein for AT1G65090.1 and .2 or 152 amino acids for AT1G65090.3) was obtained and called *sldp1-2.* For *SLDP2*, a mutant line with deletion of bases 304-379 in the first exon (resulting in frameshift and premature stop codon at position 398-400, producing a potential 107 amino acid protein) was obtained and called *sldp2-2*. For *LIPA* a mutant line with deletion of bases Δ94-214 (resulting in a frameshift and premature stop codon at position 230-232 and a potential 36 amino acid protein) was obtained and called *lipa-2*. Sequence alignments and predicted protein products of all analysed mutant lines are shown in Supplemental Data S1.

#### RNA isolation and qPCR

RNA from was isolated in triplicate from 5 mg of dry seeds using an RNA extraction kit (Monarch Total RNA Miniprep Kit, NEB). cDNA synthesis was performed with 900 ng total RNA and 100 pmol oligo(dT) primer using the Maxima Reverse Transcriptase (Thermo Scientific) according to the manufacturer’s instructions. Transcript analysis by qPCR was carried out with AT4G05320 (*POLYUBIQUITIN 10*) as reference (Czechowski *et al*., 2005). Amplification and quantification were performed with the Takyon^TM^ No Rox SYBR® MasterMix dTTP Blue Kit (Eurogentec) in the iCycler System (iQ™5 Real-Time PCR Detection System, Bio-Rad). The amplification mix contained 1x Takyon^TM^ No Rox SYBR® MasterMix dTTP Blue, 2 mM primers and 4 µl cDNA in a final reaction volume of 20 µl. The PCR program consisted of a 3 min denaturation step at 95°C followed by 40 cycles of 10 s at 95°C, 20 s at 58°C, and 40 s at 72°C.

Data analysis was performed using the 2^-ΔΔCT^ method as previously described (Livak and Schmittgen, 2001).

#### Plasmid construction

For localisation studies in pollen tubes, coding sequences of the genes of interest were cloned into pLatMVC-GW, pLatMVN-GW or pLatMCC-GW (Müller *et al*., 2017) via classical or fast Gateway® (Thermo Fisher Scientific) cloning as described before (Müller *et al*., 2017). All pLat-constructs contain a LAT52 promoter for strong expression in pollen tubes (Twell *et al*., 1991) and were verified by sequencing. A list of plasmids and primers can be found in Supplemental table S1.

For localisation studies in *N. benthamiana* leaves and Arabidopsis seedlings, cloning of pMDC32-ChC/SLDP2, encoding SLDP2 appended at its C-terminus to the red fluorescent protein mCherry (SLDP2-mCherry), pMDC32-CGFP/LIPA, encoding LIPA appended at its C-terminus to a monomerised version of GFP (LIPA-mGFP), and pMDC43/LIPA, encoding LIPA appended at its N-terminus to GFP (GFP-LIPA), was performed using Gateway cloning technology (Müller *et al*., 2017) and the binary vectors pMDC32-ChC (Kretzschmar *et al*., 2020), pMDC43 (Curtis and Grossniklaus, 2003), and pMDC32-CGFP (described below), respectively. Each binary vector contains the 35S cauliflower mosaic virus promoter and was verified by automated sequencing performed at the University of Guelph Genomics Facility.

The pMDC32-CGFP binary vector contains a Gateway recombination site followed by the full-length mGFP open reading frame, which provides for the expression of a fusion protein with a C-terminal-appended mGFP. To construct pMDC32-CGFP, the mGFP coding sequence was amplified from pRTL2/monoGFP- MCS (Shockey *et al*., 2006), using primers GFP-FP-*Pac*I (5’- CCGGCCTTAATTAAAATGAGTAAAGGAGAAGAACTTTT-3’) and GFP-RP-*Sac*I (5’- CCGGCCGAGCTCTTATTTGTATAGTTCATCCATGCC-3’), which also added 5’ *Pac*I and 3’ *Sac*I restriction sites. The resulting PCR products were digested with *Pac*I and *Sac*I and ligated into similarly-digested pMDC32-ChC to yield pMDC32-CGFP.

For yeast two-hybrid (Y2H) assays, full-length *SLDP1.3, SLDP2.1*, and *LIPA* open reading frames, as well as modified versions of the latter, were amplified from the appropriate template plasmids using PCR and primers containing the *Eco*RI and *Bam*HI restriction digest sites at the 5’ and 3’ ends, respectively. Resulting PCR amplicons were then digested with *Eco*RI and *Bam*HI and ligated into similarly-digested pGBKT7 or pGADT7, which contain the GAL4-binding domain and GAL4-activation domain, respectively (Takara Bio Inc.).

#### Particle bombardment and pollen tube microscopy

*N. tabacum* pollen tubes were transiently transformed using a gene gun. For this, 6 µg of construct DNA was coated onto approx. 0.9 mg gold particles (1 µm), shot onto freshly harvested *N. tabacum* pollen of 5 flowers per transformation. Pollen tubes were grown for 5-7 h (8-10 h for LD motility assays) in liquid pollen tube medium on a microscope slide in a humid environment (in detail methods on coating and transformation were described before (Müller *et al*., 2017). For co-transformation, 6 µg DNA of each construct were pre-mixed and then coated onto the gold particles.

For pollen tube microscopy, pollen tubes were fixed in a final concentration of 1.8 % (v/v) formaldehyde in pollen tube medium (Read *et al*., 1993) (5 % w/v sucrose, 12.5 % w/v PEG-4000, 15 mM MES-KOH pH 5.9, 1 mM CaCl2,1 mM KCl, 0.8 mM MgSO4, 0.01 % H3BO3 v/v, 30 µM CuSO4) and LDs were stained with a final concentration of 0.25 µg ml^-1^ Nile red (Sigma-Aldrich, St. Louis, Missouri, USA) or 0.5 % Lipi-Blue (Dojindo Molecular Technologies, Rockville, MD, US), as indicated. Pollen tubes prepared to monitor LD movements were stained in the same manner but no fixative was added. Micrographs were acquired as single z-sections using a Zeiss LSM 510 or a Zeiss LSM780 confocal microscope (Carl Zeiss). For excitation, 405 nm Diode was used, Lipi-Blue fluorescence was detected from 443 – 475 nm, Nile red was excited with 488 nm and detected at 583-667 nm. Constructs with mCherry were excited with 561 nm and detected at 571 – 614 nm, mVenus was excited with 488 nm and detected at 518 – 550 nm or 497-533 nm when co-imaged with Nile red. HFT 405/ 514/633-nm major beam splitter (MBS) was used.

#### CLSM-based FLIM-FRET Analysis

Confocal microscopy was performed using a Leica TCS SP8 microscope equipped with the FALCON time-correlated single photon counting system. For fluorescence-lifetime imaging of transiently transformed N. tabacum pollen tubes, images were taken with a 20×/ 0.75 objective (CS2, HC PL APO). The mVenus-tagged donor proteins and mCherry/LIPA-mCherry acceptors were excited using a pulsed white light laser operating at 514 nm or 561 nm, respectively, with a pulse rate of 40 kHz in a two channel sequential excitation mode. Fluorescence emission was detected at 525-560 nm for mVenus and 580-630 nm for mCherry using Leica HyD SMD detectors. Images were acquired until at least 100 photons per pixel were collected in the brightest channel. The format of the pictures is 512 x 512 pixels. The ROI (region of interest) selection and FLIM data fitting were performed using the LASX Single Molecule Detection software module (v3.5.5). A monoexponential reconvolution model was fitted to all decay curves for calculating the fluorescence lifetime. Donor lifetime data were exported and used for further statistical analysis and plotting in Origin 2020 (OriginLab Corp., Northampton, MA, USA).

#### Yeast two-hybrid

Directed yeast two-hybrid assays were performed according to Richardson *et al*. (2011). In brief, both bait (pGBKT7) and prey (pGADT7) vectors were co-transformed into yeast (strain PJ69-4A) using the lithium acetate transformation method (Gietz and Schiestl, 2007) and then plated on double-drop out (DDO) selection plates, consisting of synthetic dextrose (SD) media containing 2% dextrose, 0.67% yeast nitrogen base, and synthetic complete amino acid and base supplements lacking Leu and Trp (Bufferad Inc.). Selected yeast colonies were grown to log phase in liquid DDO at 30 °C and 275 rpm, then the OD_600_ was adjusted to 0.5 and 1:5 serial dilutions were carried out. 5 µl of each dilution was spotted onto both DDO and quadruple drop-out (QDO) (SD medium lacking Leu, Trp, Ade, and His) plates and grown at 30 °C for 3 days. Results shown are representative of three independent yeast transformations.

#### *N. benthamiana* infiltration and microscopy

For infiltration, leaves of 4-week-old *N. benthamiana* plants were (co)infiltrated with *Agrobacterium tumefaciens* (strain LBA4404) harbouring a selected binary vector, as described previously (Kretzschmar *et al*., 2020). All (co)infiltrations also included *A. tumefaciens* transformed with pORE04-35S::P19, which encodes the tomato bushy stunt virus gene *P19* to enhance transgene expression (Petrie *et al*., 2010).

(Co)infiltrated N. benthamiana leaves were prepared for confocal laser scanning microscopy (CLSM) by first fixing with 4% (w/v) formaldehyde, washing with 50 mM PIPES pH 7.0, and then staining with neutral lipid-specific dye monodansylpentane (MDH) (Abcepta) (Yang *et al*., 2012) at a working concentration of 0.4 mM, as described previously (Gidda *et al*., 2016). Micrographs of leaf epidermal cells were acquired as single z-sections using a Leica SP5 CLSM (Leica Microsystems) with the same excitation and emission parameters for mCherry, GFP, and MDH as reported previously (Gidda *et al*., 2016). All images of cells are representative of at least two independent experiments (i.e., infiltrations), including at least three separate (co)transformation of leaf epidermal cells.

#### Determination of 1000 seed weight and seed total fatty acid analysis

1000 seed weight was determined by manually counting replicates of 500 seeds and weighing these.

For fatty acid analysis, seeds were sieved to a size of 250 – 300 µm. Six biological replicates were performed: seeds from six different mother plants were harvested and 25 seeds each were analysed per time point and genotype, presented results are representative of two other replications of the experiment. Seeds were germinated on wet filter papers soaked in 1.6 ml H_2_O and put in a petri dish in a humid environment. Apart from seeds for dry seed analysis, seeds were stratified for 4 days at 4 °C in the dark prior to imbibition. After 4 days of stratification, 0 day samples were harvested; the other samples were placed into 16-h/8-h day/night cycle in a incubator (CU-36L/D, Percival Scientific Inc., Perry, USA) at 22 °C and a light intensity of 120 µmol m^-2^ s^-1^. Seeds and seedlings were harvested into 1 ml fatty acid methyl ester (FAME) reagent (2.5 % v/v H_2_SO_4_, 2 % v/v dimethoxypropane in methanol/toluol 2:1, v/v) (Miquel and Browse, 1992) with 30 µl of 0.33 mg/ml tri-15:0 TAG (1,2,3-tripentadecanoylglycerol ≥99%, Sigma-Aldrich, St. Louis, Missouri, USA) in toluol (ROTIPURAN® ≥ 99.5 %, Carl Roth, Karlsruhe, Deutschland) as internal standard and ground with a glass stick. Samples were then incubated at 80 °C in a water bath under constant shaking for one hour to esterify all FAs to methanol. The reaction was stopped with 1 ml of saturated NaCl-solution and vortexing. FAMEs were then extracted twice adding 1 ml of hexane, centrifuging 10 min at 2,000 x g and transferring the upper phase to a new glass tube. Hexane was evaporated and samples resuspended in 30 µl of acetonitrile (HPLC Gradient grade, Fisher Chemical, Thermo Fisher Scientific, Waltham, Massachusetts, USA). Subsequent GC-FID analysis was performed as described in (Hornung *et al*., 2002). An Agilent GC 6890 system (Agilent, Waldbronn, Germany) coupled to an FID detector equipped with a capillary HP INNOWAX column (30 m × 0.32 mm, 0.5 µm coating thickness, Agilent, Waldbronn, Germany) was used, Helium served as carrier gas (30 cm × s−1), with an injector temperature of 220 °C. The temperature gradient was 150 °C for 1 min, 150–200 °C at 15 °C min^−1^, 200–250 °C at 2 °C min^−1^, and 250 °C for 10 min. For quantification, peak integrals were determined using Agilent ChemStation for LC 3D systems (Rev. B.04.03) and used to calculate absolute amounts total fatty acids.

#### Hypocotyl measurements

All seeds used for hypocotyl analyses were sieved to a size of 250 – 300 µm prior to analyses and surface-sterilised in 6 % sodium hypochlorite solution for 15 – 20 minutes and placed on solid half-strength MS medium without sucrose (Murashige and Skoog, 1962). After a stratification for 4 days at 4 °C in the dark, seedlings were grown vertically in the light for 7 days under 16-h/8-h day/night regime or 4 h in the light and then 7 days in the dark in an incubator (CU-36L/D, Percival Scientific Inc., Perry, USA) at 22 °C and a light intensity of 120 µmol m^-2^ s-1, and hypocotyls were recorded with the Ocular scientific image acquisition software (version 1.0, Digital Optics Ltd, Auckland, New Zealand) on a binocular (Olympus SZX12 binocular, Olympus Corporation, Tokyo, Japan) attached to a camera (R6 Retiga camera, QImaging, Surrey, Canada). Hypocotyl length was measured with ImageJ software (1.52p) (Rueden *et al*., 2017) and violin plots with mean points were generated using ggplot2 package (version 3.3.2) in the R environment (version 4.0.1).

#### Seed and seedling preparation and microscopy

For time-course microscopic analyses, either seeds rehydrated for 30 min were used, or seedlings were stratified and grown on half-strength MS-medium without sucrose as described above. They were transferred to light at 07.30 am (light period 7 am – 11 pm, 16-h/8-h day/night cycle) and then analysed after 12, 24, 36 and 48 h of germination. Rehydrated seeds and seedlings were harvested into 1 ml H_2_O and directly used for microscopy after removal of seed coats. Fluorescence dyes were used at the following concentrations 1.5-6 µM Nile Red (Sigma Aldrich), 1.6 µM Bodipy 493/503, 1 µM MDY-64, or 4 µM FM4-64 (Sigma Aldrich). All stock solutions were prepared in DMSO. Seeds were additionally fixated in 1 % formaldehyde, when Z-stacks were recorded. Micrographs were taken using a Zeiss LSM780 confocal microscope (Carl Zeiss). For Nile Red excitation, 561 nm laser was used, fluorescence was detected at 571-603 nm with a 488/561 MBS. For microscopy of MDY-64 and Nile Red, laser wavelengths of 458 and 561 nm, respectively, and a 458/561 nm MBS were used, and fluorescence was detected at 463-516 and 552-631 nm, respectively. Bodipy 493/503 and mCherry were excited with a 488 and a 561 nm laser using a 488/561 MBS and detected at range of 493-568 and 586-639, respectively. eGFP and Nile Red or FM4-64 were co-excited with a 488 nm laser and a 488 MBS. Fluorescence was detected at 489-515 and 563-631 nm for eGFP and Nile Red, respectively, and 493-530 and 651-739 nm for eGFP and FM4-64, respectively.

#### Proteomic analysis

*Arabidopsis thaliana* seedlings were surface-sterilised, placed on half-strength MS-medium without sucrose, stratified for 72 h at 4 °C in the dark and then grown at 22°C under 16 h/ 8 h of light-dark regime for 38 h.

Total protein isolation of total extract (TE) and LD fractions, LD-enrichment, proteomics sample preparation including a tryptic in-gel digest, LC/MS analysis and analysis of MS/MS2 raw data was performed as previously described (Kretzschmar *et al*., 2018).

LFQ values were determined using MaxQuant software 1.6.2.10 (Cox and Mann, 2008; Cox *et al*., 2014). Perseus software (Tyanova *et al*., 2016) (version 1.6.6.2) was used for data analysis. PCA plots were created from unfiltered raw LFQ values (Supplemental Dataset S2). The libraries, the meta data file, raw data files, MaxQuant search files as well as ProteinGroup and Peptide search results created by MaxQuant are available on ProteomeXchange/PRIDE (Vizcaíno *et al*., 2014) under the identifier PXD022769.

LFQ values were normalised as ‰ of total sum of all LFQs per replicate and log_2_-transformed (rLFQ) for further analyses. For LD-enrichment analysis within one line, all proteins from TE and LD fractions together were filtered for those detected at least three times in at least one group and identified by at least two peptides (Supplemental Dataset S3).

For differential abundance analysis of LD or TE fractions between the lines, LD fraction and TE fraction were analysed separately and filtering was performed independently of the respective other fraction (Supplemental Dataset S4).

For enrichment and differential abundance analyses, rLFQ-values were imputed: missing values were replaced from normal distribution (for total extract: width 0.3, down shift 1.8; for LD fractions: width 0.5, down shift 1.8; for both fractions together: width 0.8, down shift 1.8; Supplemental Dataset S5. To obtain proteins significantly enriched on LDs, LD fractions were compared to TE fractions and analysed for proteins enriched in LD fractions. To find differentially abundant proteins between the different lines, LD fractions of mutants were compared to LD fractions of the wild type, the same was done for TE fractions. Proteins were considered LD-enriched or differentially abundant, respectively, if FDR < 0.01 and S0 > 2 (as determined by two-sided t-test with 250 randomisations). Volcano plots were created to visualise the results.

#### Electron microscopy

High pressure freezing electron microscopic analysis was performed as described before (Hillmer *et al*., 2012). Plant material was dissected from hypocotyls or cotyledons of 36 – 48 h germinated seedlings with a biopsy punch (pfm medical, Köln; 2mm diameter), submerged in freezing medium (200 mM Suc, 10 mM trehalose, and 10 mM Tris buffer, pH 6.6) transferred into planchettes (Wohlwend, Sennwald, Switzerland; type 241 and 242), and frozen in a high-pressure freezer (HPM010; Bal-Tec, Liechtenstein). Freeze substitution was performed in a Leica EM AFS2 freeze substitution unit in dry acetone supplemented with 0.3% uranyl acetate at −85°C for 16 h before gradually warming up to −50 °C over a 5-h period. After washing with 100% ethanol for 60 min, samples were stepwise infiltrated (intermediate steps of 30%, 60% HM20 in ethanol, and twice with 100% HM20 for 1h each), embedded in Lowicryl HM20 at −50 °C and polymerised for 3 d with UV light in the freeze substitution apparatus at -35 °C. Ultrathin sections were cut on a Leica Ultracut S and poststaind with 3 % aqueous uranyl acetate and lead citrate for 3 min each. Micrographs were taken at a Jeol JEM1400 TEM (Jeol Germany, Freising) equipped with a TVIPS TEMCAM F416 digital camera (TVIPS, Gauting) using EMMenue 4 (TVIPS, Gauting).

#### Bioinformatics

For sequence alignments, T-Coffee (Notredame *et al*., 2000) (http://tcoffee.crg.cat/apps/tcoffee/do:regular) was used with default settings. Sequence identity was calculated by Needleman-Wunsch global alignment of two sequences (Needleman and Wunsch, 1970) with EMBOSS needle on default settings (https://www.bioinformatics.nl/cgi-bin/emboss/needle). Helical wheel plots were created by Heliquest (Gautier *et al*., 2008) (https://heliquest.ipmc.cnrs.fr/cgi-bin/ComputParams.py) with Helix type: alpha and window size: 1_TURN. For hydrophobicity plots, ExPASy ProtScale (https://web.expasy.org/protscale/) was used with a Kyte&Doolittle scale (Kyte and Doolittle, 1982) and a window size of 9. Charge plots were created by EMBOSS explorer charge (http://www.bioinformatics.nl/cgi-bin/emboss/charge?_pref_hide_optional=0) with a window length of 5. TMDprediction was performed with ExPASy TMpred (https://embnet.vital-it.ch/software/TMPRED_form.html). Coiled-coils were predicted by ExPASy COILS (Lupas *et al*., 1991) (https://embnet.vital-it.ch/software/COILS_form.html) with a window width of 21.

#### Structure bioinformatics and modelling

Structure of LIPA (Uniprot code Q3EDG6) was downloaded from the AlphaFold2 structure database (Varadi *et al*., 2022). To calculate the structure of LIPA L80P, V98P, ColabFold with MMseq2 homology search was used (Mirdita *et al*., 2021, Preprint). Electrostatic potential was calculated using the APBS server (Jurrus *et al*., 2018).

The structure of LIPA was mapped into the Martini coarse-grain representation using the martinize2 script with the ScFix modification (Souza *et al*., 2021). The phospholipid bilayer, in total composed of 2042 phospholipid molecules, containing palmitoyl-oleoyl-phosphatidylcholine:palmitoyl-oleoyl-phosphatidylethanolamine:palmitoyl-oleoyl-phosphatidylserine:palmitoyl-oleoyl-phosphatidic acid:palmitoyl-oleoyl-phosphatidylinositol 4-phosphate:palmitoyl-oleoyl-phosphatidylinositol 4,5-bisphosphate (molecular ratio 37:37:10:10:5:1) was generated using the insane.py script (Wassenaar *et al*., 2015). MD simulations were performed with Gromacs2018 (Abraham *et al*., 2015). Lennard-Jones and electrostatic interactions were cut off at 1.1 nm, with the potentials shifted to zero at the cutoff. A relative dielectric constant of 15 was used. The neighbour list was updated every 20 steps using the Verlet neighbour search algorithm. Simulations were run in the NPT ensemble. During the production runs, the system was subject to pressure scaling to 1 bar using Parrinello-Rahman barostat with temperature scaling to 283 K using the velocity-rescaling method with coupling times of 1.0 and 12.0 ps. Semi-isotropic pressure coupling with a compressibility of 3·10^−4^ bar^−1^ was employed. Initially, the protein was placed approximately 3.0 nm away from the membrane. Subsequently, the standard MARTINI water together with Na^+^ and Cl^-^ ions at the concentration of 150 mM were added. Next, additional Na^+^ ions were added to ensure the electroneutrality of the system. The whole system was energy-minimized using the steepest descent method up to the maximum of 5000 steps and equilibrated for 10 ns with the pressure controlled by the Berendsen barostat. Production runs were performed for up to 1 μs with a time step of 20 fs. Membrane binding events were analyzed by monitoring the minimum distance between the protein and and the membrane using the gmx mindist tool in Gromacs. Membrane binding was subsequently evaluated by computing the probability density distributions using the kernel density estimation method (Srinivasan *et al*., 2021). Visualization was done using the ChimeraX and VMD program (Humphrey *et al*., 1996).

#### Accession Numbers

Sequence data from this article can be found in the GenBank/EMBL data libraries under the following accession numbers: AT1G65090 (SLDP1); AT5G36100 (SLDP2); AT1G07985 (LIPA).

### Supplemental Information

**Supplemental Table S1:** Overview of primers

**Supplementary Figure 1:** Alignment of Arabidopsis SLDP1 and SLDP2 protein isoforms

**Supplementary Figure 2:** Charge plots of SLDP

**Supplementary Figure 3:** qPCR analysis of SLDP splice variants

**Supplementary Figure 4:** qPCR analysis of SLDP mutant lines

**Supplementary Figure 5:** Images of sldp mutants.

**Supplementary Figure 6:** Time-course analysis of SLDP and LIPA mutant line LDs in hypocotyls

**Supplementary Figure 7:** Analysis of storage vacuoles

**Supplementary Figure 8:** Time-course analysis of SLDP and LIPA mutant line LDs in hypocotyls

**Supplementary Figure 9:** Transmission electron microscopy of seedlings

**Supplementary Figure 10:** Proteomic Analyses of sldp mutants.

**Supplementary Figure 11:** In silico analysis of LIPA

**Supplementary Figure 12:** Localisation analysis of LIPA in leaves.

**Supplementary Figure 13:** Co-expression analysis of SLDP and LIPA in tobacco pollen tubes.

**Supplementary Figure 14:** Coiled-coil mutants of LIPA

**Supplementary Figure 15:** Y2H assays to test for SLDP and LIPA self-interaction

**Supplementary Figure 16:** Analysis of the molecular dynamics simulations

**Supplementary Figure 17:** qPCR analysis of LIPA mutant lines

**Supplemental Data S1:** Sequence information.

**Supplemental Data S2:** Raw LFQs from proteomic analyses of LD-enriched and total extract fractions of wild-type and *sldp* mutant seedlings.

**Supplemental Data S3:** Normalised and filtered LFQs from proteomic analyses of LD-enriched and total extract fractions of wild-type and *sldp* mutant seedlings.

**Supplemental Data S4:** Imputed rLFQs from proteomic analyses of LD-enriched and total extract fractions of wild-type and *sldp* mutant seedlings.

**Supplemental Data S5:** List of LD-enriched proteins in wild-type and *sldp* mutant seedlings.

**Supplemental Data S6** Proteomic metadata table.

**Supplemental Movie S1:** LD movement in pollen tubes expressing mVenus.

**Supplemental Movie S2:** LD movement in pollen tubes expressing mVenus-LIPA.

**Supplemental Movie S3:** LD movement in pollen tubes expressing SLDP2.1- mVenus.

**Supplemental Movie S4:** LD movement in pollen tubes co-expressing mVenus-LIPA and SLDP2.1-mVenus.

**Supplemental Movie S5:** LD movement in pollen tubes expressing SLDP1.3- mVenus.

**Supplemental Movie S6:** LD movement in pollen tubes co-expressing mVenus-LIPA and SLDP1.3-mVenus.

**Supplemental Movie S7:** LD movement in pollen tubes co-expressing untagged LIPA and SLDP2.1.

**Supplemental Movie S8:** LD movement in pollen tubes co-expressing untagged LIPA and SLDP1.3

**Supplemental Movie S9:** Simulation of LIPA membrane interaction

## Acknowledgements

HEK, OV, KS, MW, GB and TI thank the German research foundation (DFG; Grants IS 273/2-2, IS 273/7-1, IS 273/10-1, IRTG 2172 PRoTECT, BR1502-15-1 and INST 186/1230-1 FUGG to Stefanie Pöggeler, INST 186/1277-1 FUGG to Volker Lipka). PS thanks the Studienstiftung des deutschen Volkes for funding. Research in RTM’s lab was supported by grants from the US Department of Energy, Office of Science, BES-Physical Biosciences program (DE-SC0016536) and the Natural Sciences and Engineering Research Council of Canada (RGPIN-2018-04629). NMD is a recipient of an Ontario Graduate Scholarship. We would like to thank Julia Matz for help in the lab, The department of Dr. Peter Rehling for granting access to their microscope and Karen Linnemannstöns for helpful discussions.

## References

Abraham, M.J., Murtola, T., Schulz, R., Páll, S., Smith, J.C., Hess, B., and Lindahl, E. (2015). GROMACS: High performance molecular simulations through multi-level parallelism from laptops to supercomputers. SoftwareX 1–2: 19–25.

Akdel, M., Pires, D. E. V, Porta Pardo, E., Jänes, J., Zalevsky, A. O., Mészáros, B., … Beltrao, P. (2021). A structural biology community assessment of AlphaFold 2 applications. BioRxiv [Preprint], 2021.09.26.461876 [accessed 11 January 2022]. Available from: https://doi.org/10.1101/2021.09.26.461876.

Bae, S., Park, J., and Kim, J.S. (2014). Cas-OFFinder: A fast and versatile algorithm that searches for potential off-target sites of Cas9 RNA-guided endonucleases. Bioinformatics 30: 1473–1475.

Baillie, A.L., Falz, A.L., Müller-Schüssele, S.J., and Sparkes, I. (2020). It Started With a Kiss: Monitoring Organelle Interactions and Identifying Membrane Contact Site Components in Plants. Front. Plant Sci. 11: 517.

Berardini, T.Z., Reiser, L., Li, D., Mezheritsky, Y., Muller, R., Strait, E., and Huala, E. (2015). The arabidopsis information resource: Making and mining the “gold standard” annotated reference plant genome. Genesis 53: 474–485.

Bewley, J. (1997). Seed Germination and Dormancy. Plant Cell 9: 1055–1066.

Bohnert, M. (2020). Tethering Fat: Tethers in Lipid Droplet Contact Sites. Contact 3: 251525642090814.

Cai, Y., Goodman, J.M., Pyc, M., Mullen, R.T., Dyer, J.M., and Chapman, K.D. (2015). Arabidopsis SEIPIN Proteins Modulate Triacylglycerol Accumulation and Influence Lipid Droplet Proliferation. Plant Cell 27: 2616–2636.

Chang, D.K., Cheng, S.F., Trivedi, V.D., and Lin, K.L. (1999). Proline affects oligomerization of a coiled coil by inducing a kink in a long helix. J. Struct. Biol. 128: 270–279.

Cheng, H.Y., Schiavone, A.P., and Smithgall, T.E. (2001). A Point Mutation in the N-Terminal Coiled-Coil Domain Releases c-Fes Tyrosine Kinase Activity and Survival Signaling in Myeloid Leukemia Cells. Mol. Cell. Biol. 21: 6170–6180.

Cheng, M.-C., Hsieh, E.-J., Chen, J.-H., Chen, H.-Y., and Lin, T.-P. (2012). Arabidopsis RGLG2, Functioning as a RING E3 Ligase, Interacts with AtERF53 and Negatively Regulates the Plant Drought Stress Response. Plant Physiol. 158: 363–375.

Cockcroft, S. and Raghu, P. (2018). Phospholipid transport protein function at organelle contact sites. Curr. Opin. Cell Biol. 53: 52–60.

Corradi, V., Sejdiu, B.I., Mesa-Galloso, H., Abdizadeh, H., Noskov, S.Y., Marrink, S.J., and Tieleman, D.P. (2019). Emerging Diversity in Lipid–Protein Interactions. Chem. Rev. 119: 5775–5848.

Cox, J., Hein, M.Y., Luber, C.A., Paron, I., Nagaraj, N., and Mann, M. (2014). Accurate proteome-wide label-free quantification by delayed normalization and maximal peptide ratio extraction, termed MaxLFQ. Mol. Cell. Proteomics 13: 2513–2526.

Cox, J. and Mann, M. (2008). MaxQuant enables high peptide identification rates, individualized p.p.b.-range mass accuracies and proteome-wide protein quantification. Nat. Biotechnol. 26: 1367–1372.

Cui, S., Hayashi, Y., Otomo, M., Mano, S., Oikawa, K., Hayashi, M., and Nishimura, M. (2016). Sucrose Production Mediated by Lipid Metabolism Suppresses the Physical Interaction of Peroxisomes and Oil Bodies during Germination of Arabidopsis thaliana. J. Biol. Chem. 291: 19734–19745.

Curtis, M.D. and Grossniklaus, U. (2003). A Gateway Cloning Vector Set for High-Throughput Functional Analysis of Genes in Planta. Plant Physiol. 133: 462–469.

Czechowski, T., Stitt, M., Altmann, T., Udvardi, M.K., and Scheible, W.R. (2005). Genome-wide identification and testing of superior reference genes for transcript normalization in arabidopsis. Plant Physiol. 139: 5–17.

Eastmond, P.J. (2006). SUGAR-DEPENDENT1 encodes a patatin domain triacylglycerol lipase that initiates storage oil breakdown in germinating Arabidopsis seeds. Plant Cell 18: 665–675.

Eisenberg-Bord, M., Shai, N., Schuldiner, M., and Bohnert, M. (2016). A Tether Is a Tether Is a Tether: Tethering at Membrane Contact Sites. Dev. Cell 39: 395–409.

Esnay, N., Dyer, J.M., Mullen, R.T., and Chapman, K.D. (2020). Lipid Droplet–Peroxisome Connections in Plants. Contact 3: 251525642090876.

Fan, J., Yu, L., and Xu, C. (2017). A Central Role for Triacylglycerol in Membrane Lipid Breakdown, Fatty Acid *β* -Oxidation, and Plant Survival under Extended Darkness. Plant Physiol. 174: 1517–1530.

Freyre, C.A.C., Rauher, P.C., Ejsing, C.S., and Klemm, R.W. (2019). MIGA2 Links Mitochondria, the ER, and Lipid Droplets and Promotes De Novo Lipogenesis in Adipocytes. Mol. Cell 76: 811–825.e14.

Gao, Q. and Goodman, J.M. (2015). The lipid droplet—a well-connected organelle. Front. Cell Dev. Biol. 3: 1–12.

Gautier, R., Douguet, D., Antonny, B., and Drin, G. (2008). HELIQUEST: A web server to screen sequences with specific α-helical properties. Bioinformatics 24: 2101–2102.

Geltinger, F., Tevini, J., Briza, P., Geiser, A., Bischof, J., Richter, K., Felder, T., and Rinnerthaler, M. (2020). The transfer of specific mitochondrial lipids and proteins to lipid droplets contributes to proteostasis upon stress and aging in the eukaryotic model system Saccharomyces cerevisiae. GeroScience 42: 19–38.

Gidda, S.K., Park, S., Pyc, M., Yurchenko, O., Cai, Y., Wu, P., Andrews, D.W., Chapman, K.D., Dyer, J.M., and Mullen, R.T. (2016). Lipid droplet-associated proteins (LDAPs) are required for the dynamic regulation of neutral lipid compartmentation in plant cells. Plant Physiol. 170: 2052–2071.

Gietz, R.D. and Schiestl, R.H. (2007). High-efficiency yeast transformation using the LiAc/SS carrier DNA/PEG method. Nat. Protoc. 2: 31–34.

Greer, M.S., Cai, Y., Gidda, S.K., Esnay, N., Kretzschmar, F.K., Seay, D., McClinchie, E., Ischebeck, T., Mullen, R.T., Dyer, J.M., and Chapman, K.D. (2020). SEIPIN Isoforms Interact with the Membrane-Tethering Protein VAP27-1 for Lipid Droplet Formation. Plant Cell 32: 2932–2950.

Hariri, H., Speer, N., Bowerman, J., Rogers, S., Fu, G., Reetz, E., Datta, S., Feathers, J.R., Ugrankar, R., Nicastro, D., and Henne, W.M. (2019). Mdm1 maintains endoplasmic reticulum homeostasis by spatially regulating lipid droplet biogenesis. J. Cell Biol. 218: 1319–1334.

Hillmer, S., Viotti, C., and Robinson, D.G. (2012). An improved procedure for low-temperature embedding of high-pressure frozen and freeze-substituted plant tissues resulting in excellent structural preservation and contrast. J. Microsc. 247: 43–47.

Hornung, E., Pernstich, C., and Feussner, I. (2002). Formation of conjugated Δ11 Δ13-double bonds by Δ12-linoleic acid (1,4)-acyl-lipid-desaturase in pomegranate seeds. Eur. J. Biochem. 269: 4852–4859.

Hugenroth, M. and Bohnert, M. (2020). Come a little bit closer! Lipid droplet-ER contact sites are getting crowded. Biochim. Biophys. Acta - Mol. Cell Res. 1867: 118603.

Humphrey, W., Dalke, A., and Schulten, K. (1996). VMD: Visual molecular dynamics. J. Mol. Graph. 14: 33–38.

Ischebeck, T., Krawczyk, H.E., Mullen, R.T., Dyer, J.M., and Chapman, K.D. (2020). Lipid droplets in plants and algae: Distribution, formation, turnover and function. Semin. Cell Dev. Biol. 108: 82–93.

Johnson, M.R., Stephenson, R.A., Ghaemmaghami, S., and Welte, M.A. (2018). Developmentally regulated H2AV buffering via dynamic sequestration to lipid droplets in Drosophila embryos. Elife 7: 1–28.

Jumper, J. et al. (2021). Highly accurate protein structure prediction with AlphaFold. Nature 596: 583–589.

Jurrus, E. et al. (2018). Improvements to the APBS biomolecular solvation software suite. Protein Sci. 27: 112–128.

Kang, B.-H. et al. (2021). A glossary of plant cell structures: current insights and future questions. Plant Cell. doi: 10.1093/plcell/koab247. Online ahead of print.

Klepikova, A. V., Kasianov, A.S., Gerasimov, E.S., Logacheva, M.D., and Penin, A.A. (2016). A high resolution map of the Arabidopsis thaliana developmental transcriptome based on RNA-seq profiling. Plant J. 88: 1058–1070.

Kory, N., Farese, R. V., and Walther, T.C. (2016). Targeting Fat: Mechanisms of Protein Localization to Lipid Droplets. Trends Cell Biol. 26: 535–546.

Kretzschmar, F.K., Doner, N.M., Krawczyk, H.E., Scholz, P., Schmitt, K., Valerius, O., Braus, G.H., Mullen, R.T., and Ischebeck, T. (2020). Identification of Low-Abundance Lipid Droplet Proteins in Seeds and Seedlings. Plant Physiol. 182: 1326–1345.

Kretzschmar, F.K., Mengel, L.A., Müller, A.O., Schmitt, K., Blersch, K.F., Valerius, O., Braus, G.H., and Ischebeck, T. (2018). PUX10 Is a Lipid Droplet-Localized Scaffold Protein That Interacts with CELL DIVISION CYCLE48 and Is Involved in the Degradation of Lipid Droplet Proteins. Plant Cell 30: 2137–2160.

Kyte, J. and Doolittle, R.F. (1982). A simple method for displaying the hydropathic character of a protein. J. Mol. Biol. 157: 105–132.

Livak, K.J. and Schmittgen, T.D. (2001). Analysis of relative gene expression data using real-time quantitative PCR and the 2-ΔΔCT method. Methods 25: 402–408.

Lundquist, P.K., Shivaiah, K.K., and Espinoza-Corral, R. (2020). Lipid droplets throughout the evolutionary tree. Prog. Lipid Res. 78: 101029.

Lupas, A., Van Dyke, M., and Stock, J. (1991). Predicting coiled coils from protein sequences. Science (80-. ). 252: 1162–1164.

Marrink, S.J., Corradi, V., Souza, P.C.T., Ingólfsson, H.I., Tieleman, D.P., and Sansom, M.S.P. (2019). Computational Modeling of Realistic Cell Membranes. Chem. Rev. 119: 6184–6226.

Mier, P., Alanis-Lobato, G., and Andrade-Navarro, M.A. (2017). Protein-protein interactions can be predicted using coiled coil co-evolution patterns. J. Theor. Biol. 412: 198–203.

Miquel, M. and Browse, J. (1992). Arabidopsis mutants deficient in polyunsaturated fatty acid synthesis: Biochemical and genetic characterization of a plant oleoyl-phosphatidylcholine desaturase. J. Biol. Chem. 267: 1502–1509.

Mirdita, M., Schütze, K., Moriwaki, Y., Heo, L., Ovchinnikov, S., & Steinegger, M. (2021). ColabFold -Making protein folding accessible to all. BioRxiv [Preprint], 2021.08.15.456425 [accessed 11 January 2022]. Available from: https://doi.org/10.1101/2021.08.15.456425.

Muliyil, S., Levet, C., Düsterhöft, S., Dulloo, I., Cowley, S.A., and Freeman, M. (2020). ADAM 17-triggered TNF signalling protects the ageing Drosophila retina from lipid droplet-mediated degeneration . EMBO J. 39:e104415.

Müller, A.O., Blersch, K.F., Gippert, A.L., and Ischebeck, T. (2017). Tobacco pollen tubes - a fast and easy tool for studying lipid droplet association of plant proteins. Plant J. 89: 1055–1064.

Müller, A.O. and Ischebeck, T. (2018). Characterization of the enzymatic activity and physiological function of the lipid droplet-associated triacylglycerol lipase AtOBL1. New Phytol. 217: 1062–1076.

Murashige, T. and Skoog, F. (1962). A Revised Medium for Rapid Growth and Bio Assays with Tobacco Tissue Cultures. Physiol. Plant. 15: 473–497.

Nakabayashi, K., Okamoto, M., Koshiba, T., Kamiya, Y., and Nambara, E. (2005). Genome-wide profiling of stored mRNA in Arabidopsis thaliana seed germination: Epigenetic and genetic regulation of transcription in seed. Plant J. 41: 697–709.

Needleman, S.B. and Wunsch, C.D. (1970). A general method applicable to the search for similarities in the amino acid sequence of two proteins. J. Mol. Biol. 48: 443–453.

Noack, L.C. and Jaillais, Y. (2020). Functions of Anionic Lipids in Plants. Annu. Rev. Plant Biol. 71: 71–102.

Notredame, C., Higgins, D.G., and Heringa, J. (2000). T-coffee: A novel method for fast and accurate multiple sequence alignment. J. Mol. Biol. 302: 205–217.

Olzmann, J.A. and Carvalho, P. (2019). Dynamics and functions of lipid droplets. Nat. Rev. Mol. Cell Biol. 20: 137–155.

Park, J., Bae, S., and Kim, J.S. (2015). Cas-Designer: A web-based tool for choice of CRISPR-Cas9 target sites. Bioinformatics 31: 4014–4016.

Petrie, J.R., Shrestha, P., Liu, Q., Mansour, M.P., Wood, C.C., Zhou, X.R., Nichols, P.D., Green, A.G., and Singh, S.P. (2010). Rapid expression of transgenes driven by seed-specific constructs in leaf tissue: DHA production. Plant Methods 6: 8.

Prinz, W.A., Toulmay, A., and Balla, T. (2020). The functional universe of membrane contact sites. Nat. Rev. Mol. Cell Biol. 21: 7–24.

Pyc, M. et al. (2021). LDIP cooperates with SEIPIN and LDAP to facilitate lipid droplet biogenesis in Arabidopsis. Plant Cell 33: 3076–3103.

Pyc, M., Cai, Y., Gidda, S.K., Yurchenko, O., Park, S., Kretzschmar, F.K., Ischebeck, T., Valerius, O., Braus, G.H., Chapman, K.D., Dyer, J.M., and Mullen, R.T. (2017). Arabidopsis lipid droplet-associated protein (LDAP) – interacting protein (LDIP) influences lipid droplet size and neutral lipid homeostasis in both leaves and seeds. Plant J. 92: 1182–1201.

Read, S.M., Clarke, A.E., and Bacic, A. (1993). Stimulation of growth of cultured Nicotiana tabacum W 38 pollen tubes by poly(ethylene glycol) and Cu(II) salts. Protoplasma 177: 1–14.

Richardson, L. G. L., Howard, A. S. M., Khuu, N., Gidda, S. K., McCartney, A., Morphy, B. J., & Mullen, R. T. (2011). Protein–Protein Interaction Network and Subcellular Localization of the Arabidopsis Thaliana ESCRT Machinery. Frontiers in Plant Science, 2: 20.

Rossini, M., Pizzo, P., and Filadi, R. (2020). Better to keep in touch: investigating inter-organelle cross-talk. FEBS J.: 288: 740–755.

Rotsch, A.H., Kopka, J., Feussner, I., and Ischebeck, T. (2017). Central metabolite and sterol profiling divides tobacco male gametophyte development and pollen tube growth into eight metabolic phases. Plant J. 92: 129–146.

Rueden, C.T., Schindelin, J., Hiner, M.C., DeZonia, B.E., Walter, A.E., Arena, E.T., and Eliceiri, K.W. (2017). ImageJ2: ImageJ for the next generation of scientific image data. BMC Bioinformatics 18: 1–26.

Rylott, E.L., Rogers, C.A., Gilday, A.D., Edgell, T., Larson, T.R., and Graham, I.A. (2003). Arabidopsis Mutants in Short- and Medium-chain Acyl-CoA Oxidase Activities Accumulate Acyl-CoAs and Reveal That Fatty Acid β-Oxidation Is Essential for Embryo Development. J. Biol. Chem. 278: 21370–21377.

Salo, V.T. et al. (2019). Seipin Facilitates Triglyceride Flow to Lipid Droplet and Counteracts Droplet Ripening via Endoplasmic Reticulum Contact. Dev. Cell 50: 478–493.e9.

Schaffer, J.E. (2003). Lipotoxicity: when tissues overeat. Curr. Opin. Lipidol. 14: 281–287.

Schapire, A.L., Voigt, B., Jasik, J., Rosado, A., Lopez-Cobollo, R., Menzel, D., Salinas, J., Mancuso, S., Valpuesta, V., Baluska, F., and Botella, M.A. (2008). Arabidopsis synaptotagmin 1 is required for the maintenance of plasma membrane integrity and cell viability. Plant Cell 20: 3374–3388.

Schauder, C.M., Wu, X., Saheki, Y., Narayanaswamy, P., Torta, F., Wenk, M.R., De Camilli, P., and Reinisch, K.M. (2014). Structure of a lipid-bound extended synaptotagmin indicates a role in lipid transfer. Nature 510: 552–555.

Schmid, M., Davison, T.S., Henz, S.R., Pape, U.J., Demar, M., Vingron, M., Schölkopf, B., Weigel, D., and Lohmann, J.U. (2005). A gene expression map of Arabidopsis thaliana development. Nat. Genet. 37: 501–506.

Schuldiner, M. and Bohnert, M. (2017). A different kind of love – lipid droplet contact sites. Biochim. Biophys. Acta - Mol. Cell Biol. Lipids 1862: 1188–1196.

Shai, N. et al. (2018). Systematic mapping of contact sites reveals tethers and a function for the peroxisome-mitochondria contact. Nat. Commun. 9: 1761.

Shai, N., Schuldiner, M., and Zalckvar, E. (2016). No peroxisome is an island - Peroxisome contact sites. Biochim. Biophys. Acta - Mol. Cell Res. 1863: 1061–1069.

Shockey, J.M., Gidda, S.K., Chapital, D.C., Kuan, J.C., Dhanoa, P.K., Bland, J.M., Rothstein, S.J., Mullen, R.T., and Dyer, J.M. (2006). Tung tree DGAT1 and DGAT2 have nonredundant functions in triacylglycerol biosynthesis and are localized to different subdomains of the endoplasmic reticulum. Plant Cell 18: 2294–2313.

Siao, W., Wang, P., Voigt, B., Hussey, P.J., and Baluska, F. (2016). Arabidopsis SYT1 maintains stability of cortical endoplasmic reticulum networks and VAP27-1-enriched endoplasmic reticulum-plasma membrane contact sites. J. Exp. Bot. 67: 6161–6171.

Souza, P.C.T. et al. (2021). Martini 3: a general purpose force field for coarse-grained molecular dynamics. Nat. Methods 18: 382–388.

Sparkes, I.A., Runions, J., Kearns, A., and Hawes, C. (2006). Rapid, transient expression of fluorescent fusion proteins in tobacco plants and generation of stably transformed plants. Nat. Protoc. 1: 2019–2025.

Srinivasan, S., Zoni, V., and Vanni, S. (2021). Estimating the accuracy of the MARTINI model towards the investigation of peripheral protein–membrane interactions. Faraday Discuss. 232: 131–148.

Sui, X., Arlt, H., Brock, K.P., Lai, Z.W., DiMaio, F., Marks, D.S., Liao, M., Farese, R. V., and Walther, T.C. (2018). Cryo–electron microscopy structure of the lipid droplet–formation protein seipin. J. Cell Biol. 217: jcb.201809067.

Thiam, A.R. and Beller, M. (2017). The why, when and how of lipid droplet diversity. J. Cell Sci. 130: 315–324.

Twell, D., Yamaguchi, J., Wing, R.A., Ushiba, J., and McCormick, S. (1991). Promoter analysis of genes that are coordinately expressed during pollen development reveals pollen-specific enhancer sequences and shared regulatory elements. Genes Dev. 5: 496–507.

Tyanova, S., Temu, T., Sinitcyn, P., Carlson, A., Hein, M.Y., Geiger, T., Mann, M., and Cox, J. (2016). The Perseus computational platform for comprehensive analysis of (prote)omics data. Nat. Methods 13: 731–740.

Ugrankar, R. et al. (2019). Drosophila Snazarus Regulates a Lipid Droplet Population at Plasma Membrane-Droplet Contacts in Adipocytes. Dev. Cell 50: 557–572.e5.

Valm, A.M., Cohen, S., Legant, W.R., Melunis, J., Hershberg, U., Wait, E., Cohen, A.R., Davidson, M.W., Betzig, E., and Lippincott-Schwartz, J. (2017). Applying systems-level spectral imaging and analysis to reveal the organelle interactome. Nature 546: 162–167.

Varadi, M. et al. (2022). AlphaFold Protein Structure Database: massively expanding the structural coverage of protein-sequence space with high-accuracy models. Nucleic Acids Res. 50: D439–D444.

Velázquez, A.P., Tatsuta, T., Ghillebert, R., Drescher, I., and Graef, M. (2016). Lipid droplet-mediated ER homeostasis regulates autophagy and cell survival during starvation. J. Cell Biol. 212: 621–631.

Vizcaíno, J.A. et al. (2014). ProteomeXchange provides globally coordinated proteomics data submission and dissemination. Nat. Biotechnol. 32: 223–226.

de Vries, J. and Ischebeck, T. (2020). Ties between Stress and Lipid Droplets Pre-date Seeds. Trends Plant Sci. 25: 1203–1214.

Waese, J. et al. (2017). ePlant: Visualizing and Exploring Multiple Levels of Data for Hypothesis Generation in Plant Biology. Plant Cell 29: 1806–1821.

Wang, Z.-P., Xing, H.-L., Dong, L., Zhang, H.-Y., Han, C.-Y., Wang, X.-C., and Chen, Q.-J. (2015). Egg cell-specific promoter-controlled CRISPR/Cas9 efficiently generates homozygous mutants for multiple target genes in Arabidopsis in a single generation. Genome Biol. 16: 144.

Wassenaar, T.A., Ingólfsson, H.I., Böckmann, R.A., Tieleman, D.P., and Marrink, S.J. (2015). Computational Lipidomics with insane : A Versatile Tool for Generating Custom Membranes for Molecular Simulations. J. Chem. Theory Comput. 11: 2144–2155.

Welte, M.A. and Gould, A.P. (2017). Lipid droplet functions beyond energy storage. Biochim. Biophys. Acta - Mol. Cell Biol. Lipids 1862: 1260–1272.

Wilfling, F. et al. (2013). Triacylglycerol synthesis enzymes mediate lipid droplet growth by relocalizing from the ER to lipid droplets. Dev. Cell 24: 384–399.

Winter, D., Vinegar, B., Nahal, H., Ammar, R., Wilson, G. V., and Provart, N.J. (2007). An “electronic fluorescent pictograph” Browser for exploring and analyzing large-scale biological data sets. PLoS One 2: 1–12.

Xie, Y., Zheng, Y., Li, H., Luo, X., He, Z., Cao, S., Shi, Y., Zhao, Q., Xue, Y., Zuo, Z., and Ren, J. (2016). GPS-Lipid: A robust tool for the prediction of multiple lipid modification sites. Sci. Rep. 6: 1–9.

Xing, H.-L., Dong, L., Wang, Z.-P., Zhang, H.-Y., Han, C.-Y., Liu, B., Wang, X.-C., and Chen, Q.-J. (2014). A CRISPR/Cas9 toolkit for multiplex genome editing in plants. BMC Plant Biol. 14: 327.

Yamazaki, T., Kawamura, Y., Minami, A., and Uemura, M. (2008). Calcium-dependent freezing tolerance in arabidopsis involves membrane resealing via synaptotagmin SYT1. Plant Cell 20: 3389–3404.

Yang, H.J., Hsu, C.L., Yang, J.Y., and Yang, W.Y. (2012). Monodansylpentane as a blue-fluorescent lipid-droplet marker for multi-color live-cell imaging. PLoS One 7.

Yang, Y. and Benning, C. (2018). Functions of triacylglycerols during plant development and stress. Curr. Opin. Biotechnol. 49: 191–198.

Yu, H., Liu, Y., Gulbranson, D.R., Paine, A., Rathore, S.S., and Shen, J. (2016). Extended synaptotagmins are Ca2+-dependent lipid transfer proteins at membrane contact sites. Proc. Natl. Acad. Sci. U. S. A. 113: 4362–4367.

Yu, J., Kang, L., Li, Y., Wu, C., Zheng, C., Liu, P., and Huang, J. (2021). RING finger protein RGLG1 and RGLG2 negatively modulate MAPKKK18 mediated drought stress tolerance in Arabidopsis. J. Integr. Plant Biol. 63: 484–493.

Zang, J., Zhang, T., Hussey, P.J., And Wang, P. (2020). Light microscopy of the endoplasmic reticulum–membrane contact sites in plants. J. Microsc. 280: 134–139.

